# Quantification of differential transcription factor activity and multiomics-based classification into activators and repressors: *diffTF*

**DOI:** 10.1101/368498

**Authors:** Ivan Berest, Christian Arnold, Armando Reyes-Palomares, Giovanni Palla, Kasper Dindler Rasmussen, Kristian Helin, Judith B. Zaugg

## Abstract

Transcription factor (TF) activity is an important read-out of cellular signalling pathways and thus to assess regulatory differences across conditions. However, current technologies lack the ability to simultaneously assess activity changes for multiple TFs and in particular to determine whether a specific TF acts globally as transcriptional repressor or activator. To this end, we introduce a widely applicable genome-wide method *diffTF* to assess differential TF activity and to classify TFs as activator or repressor (available at https://git.embl.de/grp-zaugg/diffTF). This is done by integrating any type of genome-wide chromatin accessibility data with RNA-Seq data and in-silico predicted TF binding sites. We corroborated the classification of TFs into repressors and activators by three independent analyses based on enrichments of active/repressive chromatin states, correlation of TF activity with gene expression, and activator-and repressor-specific chromatin footprints. To show the power of *diffTF*, we present two case studies: First, we applied *diffTF* in to a large ATAC-Seq/RNA-Seq dataset comparing mutated and unmutated chronic lymphocytic leukemia samples, where we identified dozens of known (40%) and potentially novel (60%) TFs that are differentially active. We were also able to classify almost half of them as either repressor and activator. Second, we applied *diffTF* to a small ATAC-Seq/RNA-Seq data set comparing two cell types along the hematopoietic differentiation trajectory (multipotent progenitors – MPP – versus granulocyte-macrophage progenitors – GMP). Here we identified the known drivers of differentiation and found that the majority of the differentially active TFs are transcriptional activators. Overall, *diffTF* was able to recover the known TFs in both case studies, additionally identified TFs that have been less well characterized in the given condition, and provides a classification of the TFs into transcriptional activators and repressors.

## INTRODUCTION

Transcription factors (TFs) are important for orchestrating coordinated and dynamic responses to intra-and extracellular stimuli and regulating a multitude of biological processes. Indeed, since many signaling cascades end in the activation of a particular set of TFs, observing a change in overall TF activity can serve as a good read-out of signaling pathways that regulate them (Kim et al., 2007). Transcriptional regulation is largely influenced by cell type specific features such as cofactors, cooperative binding partners and local chromatin environment (Whyte et al., 2013). Adding to this complexity, many TFs can act as transcriptional activators and repressors depending on the cell type and growth condition (Han et al., 2015, 2018). Thus, to correctly interpret the downstream effects of a change in abundance of a given TF, it is important to understand its global mode of action within the specific context of the study.

TFs are typically lowly abundant proteins, which makes it difficult to detect them in proteomics experiments (Kim et al., 2007; Teng et al., 2008), and even if they can be detected, their abundance and activity do not necessarily correspond since TFs are highly regulated at the post-translational level. On the other hand, chromatin immunoprecipitation followed by sequencing (ChIP-Seq), which is the gold-standard technique for measuring genomic TF binding events, provides information only for one TF at a time and does not detect global changes in TF activity unless specific experimental normalisation methods are used (e.g. spike-ins (Bonhoure et al., 2014)). Neither proteomics nor ChIP-Seq experiments can give any insights into their mode of action. Finally, luciferase assays can measure the activity and mode of action for a specific TF at a specific location and are therefore fairly low throughput (Komatsu et al., 2010; Liu et al., 2011). Databases like *TRRUST* (Han et al., 2015, 2018) collect annotations of regulation modes of TFs based on literature text mining and provide a comprehensive resource for well-studied TF-target interactions. However, for the vast majority of TFs, there is no consensus about their molecular functional mode of action. A general framework for determining differential activity of TFs between conditions and classifying TFs into transcriptional activators and repressors in a cell-type and condition-specific manner is currently lacking.

Towards closing these gaps, we have developed an approach called *diffTF* to estimate global changes in TF activities across conditions or cell types, and classify TFs into activators and repressors based on the integration of genome-wide chromatin accessibility or histone mark ChIP-Seq data with predicted TF binding sites and RNA-Seq data. We corroborated the classification of TFs into repressors and activators by three independent analyses. First, we showed that repressors and activators were enriched in repressive and active chromatin states, respectively. Second, we confirmed that expression levels of repressors were anti-correlated with their target genes while they were positively correlated with their activators. Third, we obtained activator-and repressor-specific chromatin footprints based on TFs with a known mode of action, and found that this agreed very well with the footprints obtained from the factors as classified by *diffTF*.

We applied this approach to two case studies, one comparing two patient cohorts each with a large number of heterogeneous samples, the second comparing two cell types along a differentiation trajectory each with a small number of homogeneous samples. For the first study, we obtained a large ATAC-Seq dataset of chronic lymphocytic leukemia (CLL) samples from Rendeiro et al. ((Rendeiro et al., 2016)) from > 50 patients and a total of over 1 billion reads and show that the quantification of differential TF activity by *diffTF* is highly robust with respect to a wide range of parameter settings. We recapitulate many known TFs associated with CLL and propose several novel TFs that are involved in processes related to CLL biology such as the circadian clock. Furthermore, were were able to classify ~40% of these TFs (186) into activators and repressors, thus reconciling some biological processes that seem driven by activators and repressors at the same time. For the second case study we performed ATAC-and RNA-Seq on murine multipotent progenitors (MPP) versus granulocyte-macrophage progenitors (GMP) in quadruplicate. Again, with *diffTF* we were able to identify the known driver TFs of the differentiation process, and we found that the majority of the highly differentially active TFs are acting as transcriptional activators.

Finally, the approach has been successfully applied to identify TFs that are specifically associated with TET2, an enzyme involved in DNA demethylation (Rasmussen et al., 2018), and to identify novel driving factors in pulmonary artery hypertension (Reyes-Palomares et al., in preparation).

## RESULTS

### Conceptual derivation of using open chromatin as read-out of differential TF activity

We define TF activity as the effect of a TF on the state of chromatin as measured by chromatin accessibility assays (e.g. ATAC-Seq, DNase-Seq) or ChIP-Seq for active chromatin marks (e.g. H3K27ac). This definition is based on our earlier work where we showed that genetic variants affecting H3K27ac signal across individuals (hQTLs) can be explained by disruptions of TF motifs whenever the hQTL-SNP overlaps with a TFBS (Grubert et al., 2015). Even though the exact mechanisms of how changes in TF affinity translate to the chromatin level are unknown, TF activity likely plays a causal role in mediating the effect of the DNA variant onto chromatin marks (Liu et al., 2015). By reversing this argument, we propose to use the aggregate changes in chromatin accessibility in the vicinity of putative binding sites of a TF as a read-out for its change in activity (**Suppl. Fig. 1**). A similar concept has been proposed in other tools that estimate TF activity based on ATAC or DHS data (Baek et al., 2017); (Schep et al., 2017).

Based on this concept, we developed *diffTF*, which is a computational approach to globally assess differential TF activity between two conditions (basic mode; **Fig. 1a, Suppl. Fig. 2**) and classify TFs into activators and repressors (classification mode, see below; **Fig. 1b**). It is based on any data that measures active/open chromatin, putative binding sites for TFs of interest, and optionally, for the classification mode only, matched RNA-Seq data. Briefly, for the basic mode, it requires *in silico* TFBS that can be obtained using position weight matrices (PWMs) from a database such as *HOCOMOCO* (Kulakovskiy et al., 2013) *and a PWM scanning algorithm such as PWMScan* (Ambrosini et al., 2018) for all TFs, or from a database of ChIP-Seq data, such as *ReMap* (Griffon et al., 2015). For each TFBS it then calculates the difference between two conditions and summarizes their change in accessibility across all binding sites of a given TF. In this step it also normalises for the GC content of the respective TFBS (**Suppl. Fig. 3**). The significance is assessed using either an empirical or analytical procedure. The former assesses the significance of the differential TF activity by comparing the real data against the distribution of values obtained from permuting the sample labels. The analytical procedure, which is particularly useful if the number of samples is too low for performing a reasonable number of permutations, calculates the p-value explicitly based on a t-statistic and estimated variance (see Methods for details and guidelines and **Suppl. Fig. 4**). In the basic mode, *diffTF* outputs differential activity and p-value for each TF, which together allow the identification of a set of TFs that show a significantly higher activity in one of the conditions (**Fig. 1a**).

**Figure 1.**
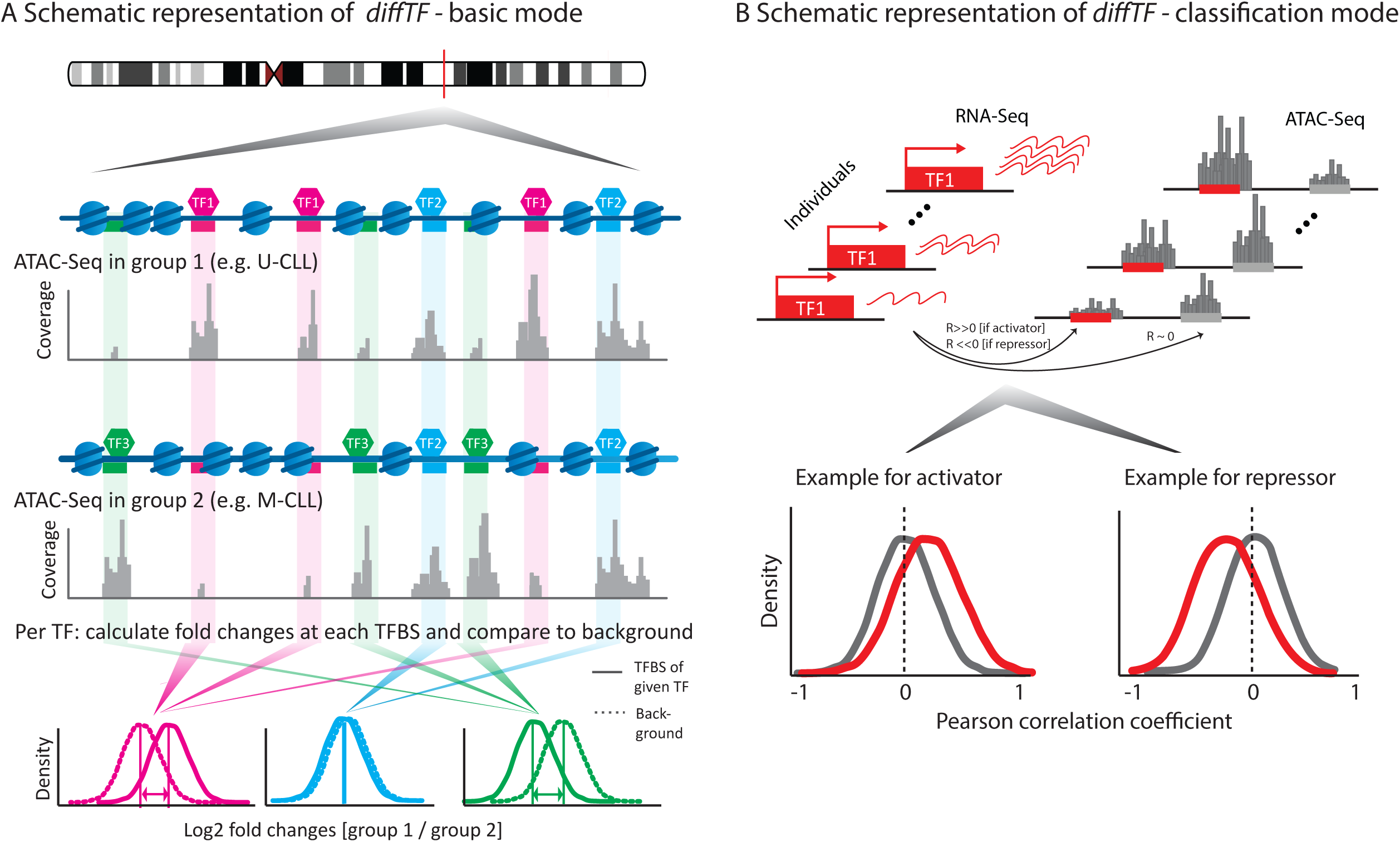
Schematic representation of the diffTF workflow. **(a)** A simplified workflow illustrates the principle upon which *diffTF* is based: it calculates a fold-change between two conditions for each binding site of a given TF and compares this distribution to a background set of fold-changes obtained from GC-content matched loci that do not contain the putative TFBS. The difference in distribution is assessed in significance and effect size and visualized in a volcano plot where the y-axis indicates statistical significance and the x-axis shows the effect size. (For a detailed workflow see **Suppl. Fig. 1** and **2**). **(b)** *Schematic representation of the classification approach*: correlation of TF expression level with the accessibility of its target sites. If the distribution of correlations between a TF’s RNA-level and the chromatin accessibility at its target sites is more positive than the background distribution (accessibility at non-target sites), the TF is classified as an activator in the particular biological environment; if negative, it is classified as a repressor. Correlations close to 0 are classified as undetermined.

### Conceptual derivation of using open chromatin and RNA to classify TFs as transcriptional activators and repressors

Surprisingly, little is known about whether a certain TF acts mostly as transcriptional repressor or activator, and based on literature text mining, most TFs have been annotated multiple times as both activator and repressor (Han et al., 2018) (**Suppl. Fig. 12b**) (Han et al., 2018) indicates that the cell type or other external factors are important in determining a TF’s main mode of action. Therefore, we devised a cell-type specific and data-driven, multiomics approach to classify TFs into activators and repressors within the framework of *diffTF* that can be run on top of the basic mode (classification mode). Our classification framework is based on the fact that increasing abundance of an activator TF results in increased transcription of its target genes (and vice versa for repressors). Yet transcription is difficult to measure since a typical RNA-Seq experiment measures the steady-state RNA level regulated by transcription and degradation. We reasoned that measures of chromatin activity (such as accessibility) is a more direct read-out for the mode of action of TFs. Based on this, we implemented an activator/repressor classification scheme in *diffTF* using RNA-Seq data as an estimate for TF abundance. For each TF, we calculate the correlations across individuals between its expression level and the ATAC-Seq signal in its putative target peak (**Fig. 1b**). Each TF is then classified (i) as an activator when it shows an overall positive correlation with the ATAC-Seq signal at its putative target sites, or (ii) as a repressor for an overall negative correlation, or (iii) as undetermined if the distribution of correlations is not significantly different from the peaks that did not overlap its putative binding sites (see also Methods and **Suppl. Fig. 12a**). The assumptions underlying this classification are tested in the context of case-study I (see below).

### Case-study I: Quantify differential TF activity in a large ATAC-Seq dataset in CLL

We sought to apply *diffTF* to a large ATAC-Seq data set in a biological setting that is well-studied so that we could assess the technical robustness and its power to recover relevant biological signal. To do so, we identified a large ATAC-Seq data-set comparing different subtypes of the extensively studied cancer chronic lymphocytic leukemia (CLL) (Rendeiro et al., 2016) as an ideal dataset.

Chronic lymphocytic leukemia (CLL) is one of the most frequent types of cancer in the Western world, particularly among the elderly. There are two major subtypes of CLL, which are defined based on the mutation status of the IgHV locus (mutated: M-CLL and unmutated: U-CLL). In M-CLL, B-cells go through normal affinity maturation with the aid of T-helper cells and undergo multiple rounds of IgHV somatic hypermutation to produce high affinity B-cell receptors (BCR). This process is essential for their development, survival and growth (Neu and Wilson, 2016). In contrast, U-CLL B-cells reach their affinity maturation point in an unregulated manner, and without involvement of T-helpers (Chiorazzi and Ferrarini, 2011). Overall, this leads to worse clinical outcomes such as shorter survival time and higher frequency of relapse after treatment (Furman et al., 2014).

The CLL dataset is comprised of ATAC-Seq data for a cohort of 88 CLL patients stratified by the mutation status of the IgHV locus (34 U-CLL, 50 M-CLL, 4 unclassified). After data processing and quality control, 25 and 27 U-CLL and M-CLL samples remained, and we applied *diffTF* in the basic mode to identify the differences in TF activities between U-CLL and M-CLL (**Suppl. Fig. 5-7**, see Supplementary Methods for more details).

In total we identified 67 TFs that are differentially active (FDR < 10%) between the two subtypes (**Fig. 2a**; **Suppl. Table 1**). About ~41% of the differentially active TFs have previously been associated with CLL and mostly (90%) agree with the reported condition (i.e., mutated or unmutated; **Suppl. Table 2**), thus providing a strong biological validation of the approach. The remaining 59% may represent novel candidate TFs that can advance our understanding of the disease etiology in general and the biological differences of mutated and unmutated patients in particular.

**Figure 2.**
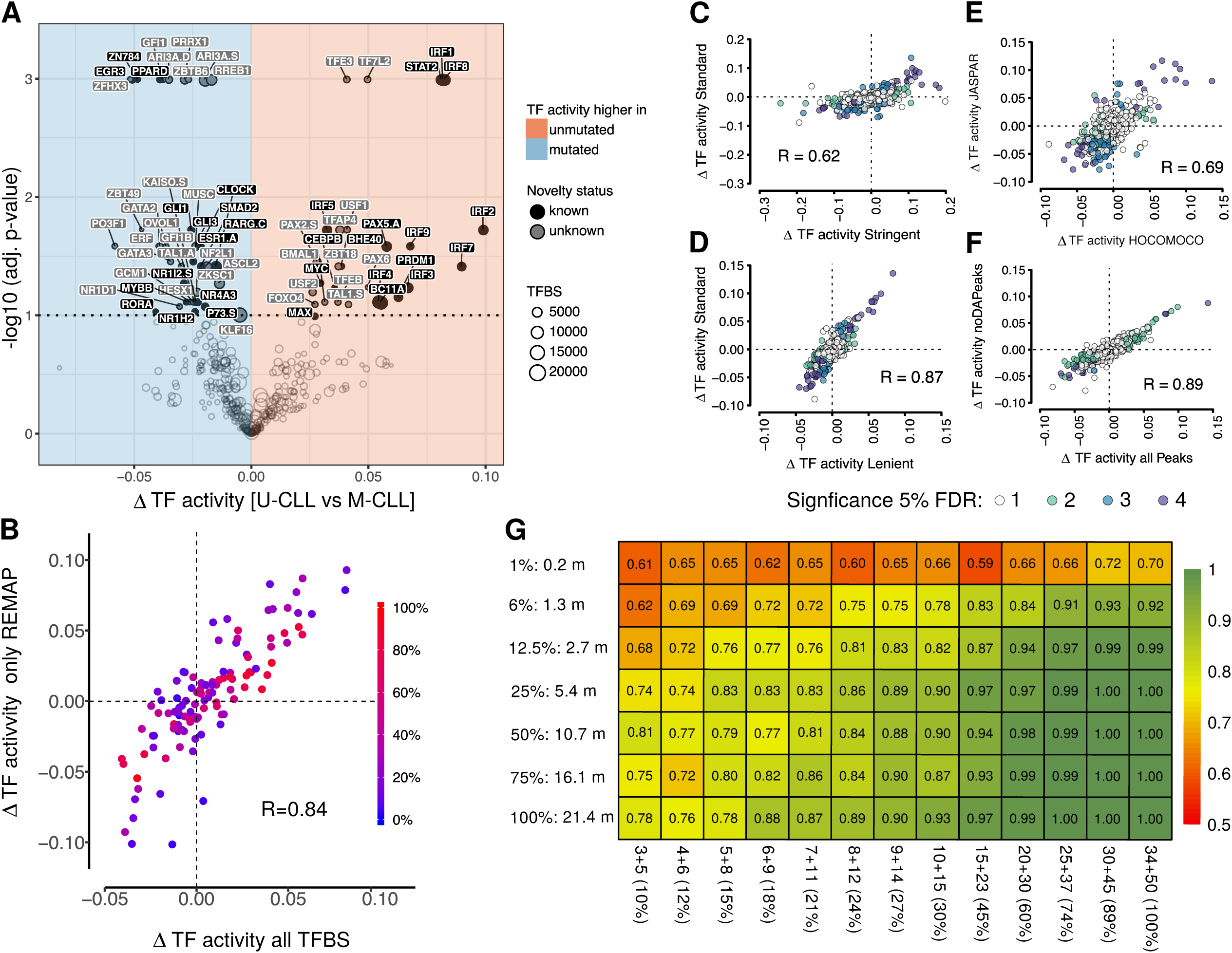
diffTF results for the CLL dataset, experimental validation and technical robustness of the method. **(a)** Volcano plot of differential TF activity between U-CLL (n=27) and M-CLL (n=25) patients. The y-axis denotes statistical significance (-log10 transformed). TFs that pass the significance threshold (5% FDR; dotted line) are labeled and colored according to their novelty status (see text and **Suppl. Table. 2**). “#TFBS” denotes the number of predicted TF binding sites in the peak regions for this analysis. (b)-(f): Technical robustness of *diffTF*. Scatterplots of the differential TF activity from all TFs for two different *diffTF* analyses are shown. Each point represents one TF. For (c-f), colors represent significance at 5% FDR (white– not significant in either analysis; light green and light blue – significant for the analysis on the x-axis or y-axis, respectively; purple – significant for both analyses). **(b)** Comparison of all predicted TFBS and TFBS experimentally validated by ChIP-Seq data from ReMap. See also **Suppl. Fig. 8**. **(c-d)** Comparison for different p-value thresholds in *PWMScan* to predict TFBS:(c) standard vs. stringent (i.e., 1e-5 vs. 1e-6) and (d) standard vs. lenient (i.e., 1e-5 vs. 5e-5) for a total of 628 TFs for which binding sites were retrieved for both scanning modes. **(e)** Comparison of *diffTF* results based on *HOCOMOCO* v10 vs. *JASPAR* 2018 as input for the 412 TFs for which a motif was available in both databases. **(f)** Comparison of the full consensus peak set (*allPeaks*) and only the non-differentially accessible peaks (*noDApeaks*; n=640 TFs from *HOCOMOCO*). **(g)** Robustness analysis based on sequencing depth and sample size. Each cell in the heatmap shows the fraction of TFs that showed the same direction of change as in the full dataset for varying degrees of down-sampling sequencing depth and number of samples, (5% FDR), averaged over 50 independent repetitions to minimize sampling noise. Only TFs that were deemed significant in the full dataset are considered (see also **Suppl. Fig. 11**). Sequencing depth is shown as a fraction of the original data and median number of reads across samples, while the number of samples is given as unmutated + mutated.

Before focusing on the biological interpretation of the specific TFs, we used this dataset to assess the technical robustness of *diffTF* with respect to TF binding site predictions. First, we compared the results of *diffTF* when using putative vs. ChIP-Seq validated TF binding sites since predicting TF binding sites is inherently noisy and may result in many false positive sites when compared to ChIP-seq experiments (Landt et al., 2012). We observed a very strong correlation of the resulting TF activity differences (r=0.81; **Fig. 2b**, **Suppl. Fig. 8**), which indicates that *diffTF* is robust with respect to false positive binding sites. Second, we assessed the parameters for TF binding site predictions and found that neither the nucleotide composition of the predicted binding sites for *PWMScan* (**Suppl. Fig. 9**) nor the motif database (*JASPAR* vs. *HOCOMOCO*) had a strong impact on the differential TF activity (r=0.87, 0.62 and 0.69, respectively, **Fig. 2c-e**). Third, we assessed whether the size of the region surrounding the TF binding site from where signal was extracted (ranging from just the 7-25 bp long binding site to additional 600 bp upstream and downstream) had an impact on the results. The resulting differential TF activities were strongly correlated (r>0.9 for 50-600 bp and r=0.76 for the binding sites only; see Supplement and **Suppl. Fig. 10**). Additional robustness tests are described in the supplement.

We also assessed the potential of *diffTF* to detect differential TF activities in experiments with little biological signal. For this, we removed high-signal regions (i.e. differentially accessible peaks at 5% FDR; see Methods) and compared the resulting differential TF activities to those of the full set. We found that they were very similar for both sets (r=0.89), thus demonstrating the power of *diffTF* to capture the differential TF activities by summarising the subtle changes in chromatin accessibility across many TFBS genome-wide (**Fig. 2f**).

Finally, we assessed the dependency of *diffTF* results on sample size and sequencing depth. Intriguingly, we found the results highly congruent across a wide range of sample sizes and sequencing depths, with the majority of the significant TFs from the full dataset changing in the same direction in the subsampled data (**Fig. 2g**, **Suppl. Fig. 11**). Generally, the number of samples appears more important than read depth, and results using the full set were consistent for a coverage as low as 1-5 million processed reads per sample (see Methods). Although the subsampling results are dataset-specific and difficult to generalize they provide guidelines for the applicability of *diffTF* and are in line with single-cell ATAC-Seq data analysis that also show robustness for low coverage at the level of genome-wide summary statistics (Mezger et al., 2018).

In summary, these results establish the robustness of *diffTF* in quantifying differences of TF activities, and demonstrate that aggregating signal across all binding sites is a powerful and sensible approach to overcome limitations such as low coverage and little underlying biological signal.

### *diffTF* proposes many novel TF candidates

We next focused on the biological interpretation of the differentially active TFs (FDR 10%) between M-CLL and U-CLL patients (**Fig. 2a**; **Suppl. Table 1**). Since TF binding motifs, which are the basis of *diffTF* (and any other tool that is based on predicted TF binding sites), can be very similar between TFs of the same family, we decided to group TFs into TF-motif families using the PWM clustering tool *Rsat* (Medina-Rivera et al., 2015); clusters available at https://bit.ly/2J9TaaK). The resulting clusters showed overall consistent activity changes within a TF family (**Fig. 3a**), with the exception of cluster 17 that can be explained by a prominent split into two branches early in the clustering into NFAT and NF?B factors, which show more activity in U-CLL and M-CLL, respectively (**Fig. 3c**).

**Figure 3.**
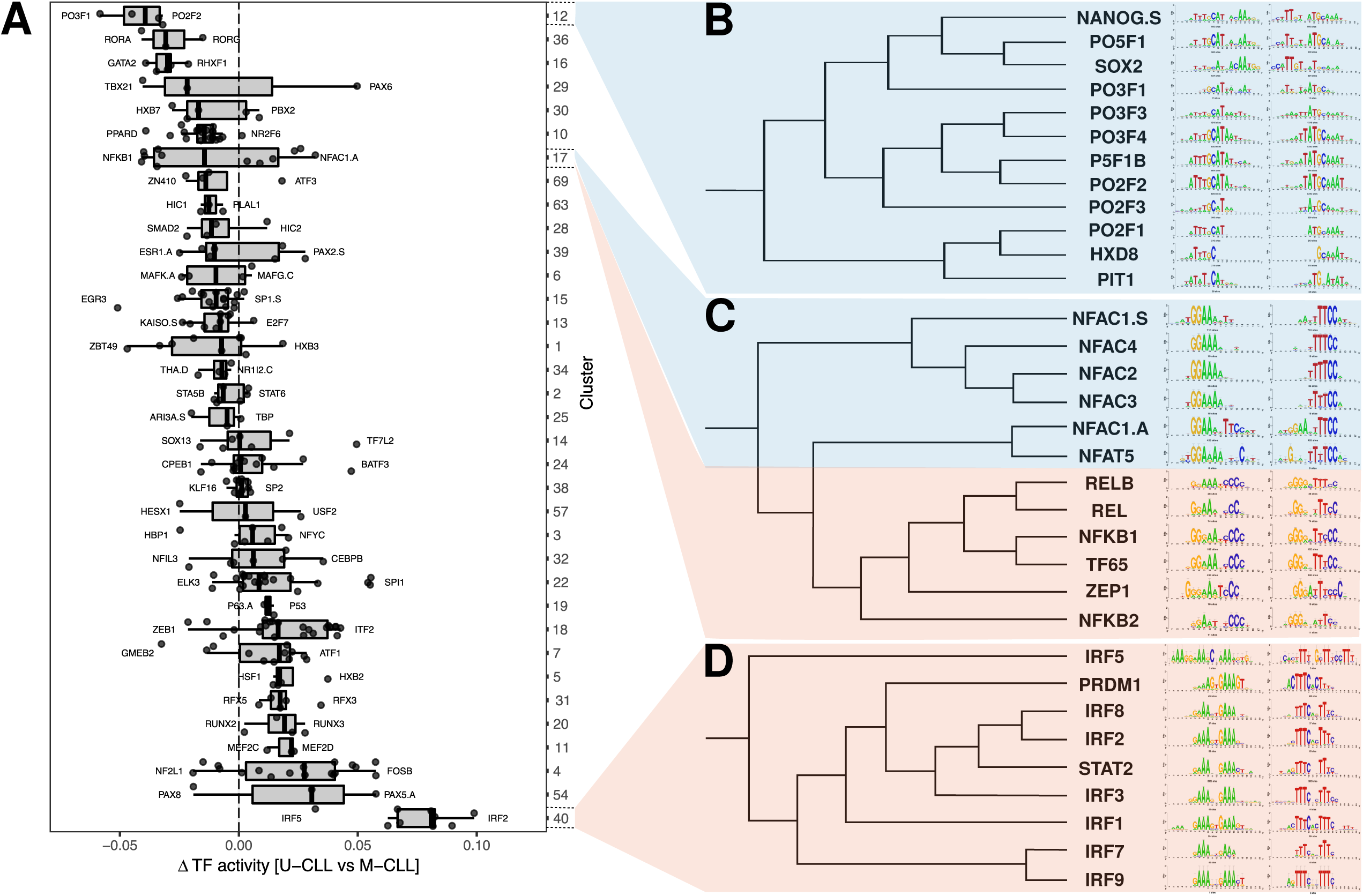
Clustering of TFs based on the similarity of their PWMs. **(a)** Boxplot of PWM clusters with at least 3 members as defined by RSAT for the differential TF activity between U-CLL and M-CLL. In each cluster, the most negative and most positive TF is labeled. (b)-(d): RSAT clustering output and tree for specific clusters. **(b)** Cluster 12 (POU family), the most distinct cluster for M-CLL patients. **(c)** Cluster 17, which has two distinct subclusters, representing the NFAT and NFKB family, respectively. **(d)** Cluster 40 (IRF family), the most distinct cluster for U-CLL patients.

The most active TF cluster in U-CLL is the IRF family and STAT2 (cluster 40; **Fig. 3d**), both of which have been associated with disease onset and progression, and harbour several CLL susceptibility loci (Arvaniti et al., 2011; Havelange et al., 2011; Slager et al., 2013). It is followed by the PAX TFs (cluster 54), which affect B-cell to plasma cell differentiation (PAX5), that is linked with cell survival and poor prognosis in CLL (Ghamlouch et al., 2015). Another prominent set of regulators are the members of the AP-1 complex (cluster 4), which increase proliferation and play an important role in driving the invasive nature of U-CLL (Mittal et al., 2013). Finally, we found c-MYC, which is involved in cell proliferation and differentiation and is highly abundant in U-CLL (Landau et al., 2015; Yeomans et al., 2016).

For M-CLL, we identified TFs that regulate and possibly reduce apoptosis, regulate cell cycle and suggest normal functionality of B-cells through the classical BCR, NF-kB and Wnt signaling pathways. The most active TF family in M-CLL patients is that of the POU TFs, also known as Oct factors (cluster 12; **Fig 3b**), which regulate B-cell development and immunoglobulin production, therefore promoting survival of the lymphoma cells (Heckman et al., 2006). This is followed by the ROR factors (cluster 36), which together with Wnt5a activate NF-kB-dependent survival signaling in CLL (Minami et al., 2010), and the GATA family (cluster 16), which is known to prime HSCs towards the lymphoid lineage and increase self-renewal of the stem cells in CLL (Kikushige et al., 2011). Other examples include the EGR family, whose motifs are enriched in aberrantly hypomethylated CpG sites in CLL (Oakes et al., 2016), PPARD, which has recently been linked to M-CLL through its effect on metabolic pathways in cancer cells (Li et al., 2017), and members of the GLI family, which are part of the Hedgehog signaling pathway and regulate apoptosis, thereby supporting survival of M-CLL cells (Kern et al., 2015).

Among the novel factors associated with U-CLL, we found several TFs (i.e. BMAL1, CLOCK, and NR1D1) that are involved in the regulation of the circadian clock, which has recently been proposed as hallmark of cancer (El-Athman and Relógio, 2018). Moreover, we found members of the basic helix-loop-helix family, such as BHE40, a regulator of mitotic division (D’Annibale et al., 2014), which is essential for the development of B1-a cells (Kreslavsky et al., 2017) and TFAP4, TFE3 and TFEB, for which there are known cases of gene-fusions in renal cell carcinoma (Kauffman et al., 2014). Another set of novel TFs more active in M-CLL are associated with pathways relevant for cancer-and B-cells such as escape from apoptosis (ZN784) (Kasim et al., 2017), regulation of cell cycle progression (ZBTB6) (Chevrier et al., 2014), and selection of B-cells and promotion of fetal B lymphopoesis (ARID3A) (Zhou et al., 2015). The GFI1 family (cluster 35) is less active in U-CLL and their expression and activation might influence and decrease rates of apoptosis in B-cells (Coscia et al., 2011).

In summary, these results show that *diffTF* is able to recapitulate much of the known biology of the two subtypes of CLL and, in addition, identifies several more factors that are likely to be implicated in the disease.

### Determination of the molecular function of TFs: transcriptional repressors and activators

The paragraphs above have shown that *diffTF* can identify TFs that alter their activity across different types of CLL patients. However, to gain mechanistic insights into some of the regulatory differences between U-CLL and M-CLL, it is crucial to know whether a TF acts as activator, in which case a higher abundance would generally result in increased transcription of its target genes, or repressor, in which case an increase in abundance would be accompanied by decreased target gene transcription (**Fig. 1b**). To do so, we employed the classification mode of *diffTF*, which integrates the ATAC-Seq data with RNA-Seq to classify TFs as activators or repressors. For this, we first needed to test the global assumption that repressors and activators have an opposing effect on chromatin accessibility that underlies our classification framework. For activators, the expectation is that an increased TF abundance will increase the accessibility at its targets sites. For a repressor, however, the relationship between abundance and accessibility at its binding site is less straightforward: On the one hand the binding of the factor itself will increase the accessibility locally, while on the other hand, repression is globally associated with closed chromatin. To understand the effect of repressors and activators on chromatin accessibility and derive general principles, we compared the accessibility footprint (Tn5 insertion sites) of a well-known repressor (REST) and a well-known activator (STAT2) that are active in our cell type. We observed that for the repressor REST, there is indeed an increase in accessibility at its motif, which likely reflects the binding of the TF itself. Importantly, however, the accessibility drops to below the genome-wide average within 10 bp from the center of the motif (**Fig. 4a, bottom**). In contrast, for the activator STAT2, we observed increased chromatin accessibility outside its core binding site, which only slowly drops to the genome-wide average over a distance of >100 bp from the center of the motif, likely representing the effect of the TF on opening the surrounding chromatin (**Fig. 4a, top**). This shows that, while there is an increase in accessibility for repressors at the immediate binding site, the surrounding chromatin is highly compact while it is open for the activator. This is in line with a previous observation on EGR and SP4 (Baek et al., 2017) and justifies our classification approach implemented in *diffTF.*

**Figure 4.**
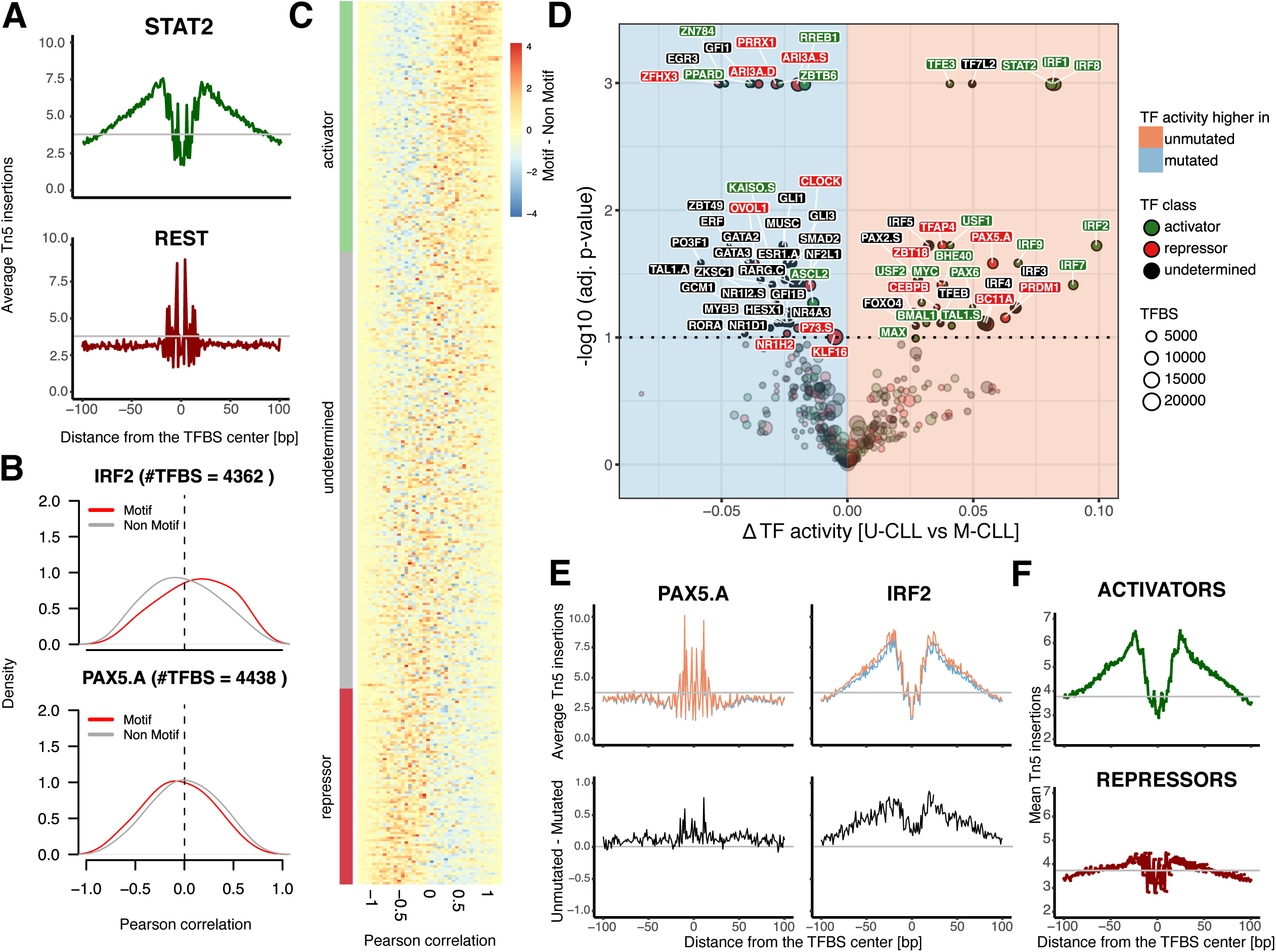
Classification of TFs into activator or repressor based on RNA-Seq and ATAC-Seq data. **(a)** Exemplary footprints for a well-known activator (STAT2, top) and repressor (REST, bottom). The x-axis depicts the distance in bp from the TFBS center, the y-axis denotes the number of average Tn5 insertions, normalised to the library size and numbers of samples across U-CLL and M-CLL. TFBS were predicted by *PWMscan* and only those overlapping with open chromatin have been considered. The solid black line indicates the average insertion sites within accessible chromatin. **(b)** Distributions of the Pearson correlations between TF expression and ATAC-Seq signal at all putative TFBS (red line), and background distribution of TFBS not containing the motif of interest (grey line) for two specific TFs, IRF2 (top) and PAX5 (bottom). **(c)** Summary of (b) across all TFs in the form of a heatmap showing the differences in Pearson correlation of putative target peaks and background for each TF (i.e., subtracting the black from the red line in (b)). Each TF is one row and is annotated as activator (green), undetermined (grey) or repressor (red) as classified by *diffTF*. **(d)** Same as Fig. 2a, but with the TFs labeled with their predicted role as activator (green), undetermined (black) or repressor (red). See Fig. 2a for details. **(e)** Footprint analysis for an activator (IRF2, right) and repressor (PAX5.A, left) as classified by *diffTF*. The top row shows the footprints separately for M-CLL and U-CLL (blue and orange, respectively) based on the normalized number of Tn5 insertions, while the bottom row highlights their differences (U-CLL - M-CLL) are shown. See (a) for axis descriptions. **(f)** Summary footprint for all activators (top, green) and repressors (bottom, red). See (a) for axis descriptions.

Applying this reasoning to the CLL dataset, where RNA-Seq data was available for eight individuals (after QC; see supplement), we were able to classify 44% of the expressed TFs as either activators or repressors (**Fig 4b-f**; n=186). Among the top activators are the IRFs, which are well known transcriptional activators (Yanai et al., 2012) and which showed the same footprint pattern as STAT2 (**Fig. 4e**). Among the top repressors, we found PAX5, which has been shown to repress the activity of BLIMP-1 (Yasuda et al., 2012) and also shows a footprint similar to the repressor REST (**Fig. 4b,e**). To assess the binding properties of activators and repressors globally we performed an unbiased footprinting analysis for all TFs deemed significant in CLL. Importantly, we found that the aggregate signal across all repressors produced a footprint similar to that of REST, while the footprint for activators looks similar to STAT2, again indicating that repressors and activators have very distinct open chromatin footprints (**Fig. 4f**). Clustering of the footprints of the individual TFs revealed four major classes. Class I is characterized by low levels of Tn5 insertions in the motif and high levels in the adjacent regions and its members are mainly classified as activators. Class II comprises of TFs with very low accessibility overall (mainly repressors). Class III contains TFs with high accessibility at the binding site and low accessibility in the adjacent regions, mainly classified as repressors but including a few activators. Finally, Class IV comprises TFs with a footprint that neither resembles an activator nor a repressor (**Suppl. Fig. 13**). This clustering indicates that TFs with a Class I footprint are likely classified as activators. In contrast, Class III footprints (like REST) are more likely classified as repressor, even though there might be some activator TFs that with a similar footprint. Overall, it seems that TF footprints correlate well with the molecular mode of action of a TF as identified by *diffTF*.

When investigating TF families as defined above with the RSAT clusters (**Fig. 3**), we found that TFs from the same PWM cluster are often classified both as activators and repressors (**Suppl. Fig. 14**), supporting the hypothesis that the molecular function of a TF is highly variable. The exceptions are cluster 40, containing mainly members of the IRF family, and cluster 17 that contains both NFAT and NFKB TFs, which are mostly classified as activators. The circadian regulators provide an example of why it is important to know the mode of action of a particular TF: When analysing the differential TF activities, it appears as if BMAL1 is more active in M-CLL while the other two TFs (CLOCK and NR1D1) are more active in U-CLL (**Fig 4d**). However, since BMAL1 is an activator, while CLOCK and NR1D1 are repressors, all three circadian factors are consistently more active in M-CLL, albeit with a contrary effect on their target genes.

To assess the validity of our repressor/activator classification, we chose three independent approaches. First, we used *chromHMM* chromatin states for primary B-cells from the Epigenomic Roadmap (Roadmap Epigenomics Consortium et al., 2015) to assess whether activators and repressors are preferentially located in active and repressive states, respectively. Indeed, we observed that the fraction of TFBS in active chromatin states was significantly larger for activators than for repressors, and vice versa for heterochromatin and repressive states (**Fig. 5a**, see also **Suppl. Fig. 15**), thus corroborating our classification of their molecular mode of action. Second, we assessed whether the direction in gene expression changes of TFs between U-CLL and M-CLL was in agreement with their differential activity and molecular mode of action. Again, our observations were in line with our expectations: activators showed a positive correlation of activity and expression change (r=0.19, P=0.05) while repressors showed a negative relationship (r=-0.32, P=0.0033; **Fig. 5b**). Third, we checked whether the expression of target genes of a given TF changes in the same direction as its activity calculated by *diffTF*, regardless of the TF’s classification as activator or repressor, and again found that this was the case for both activators and repressors (**Fig. 5c,** see Methods). In summary, these observations provide three independent lines of evidence that our approach implemented in *diffTF* is able to classify TFs globally by their mode of action. The fact that the correlations are in the expected direction but not perfect are likely reflecting that TFs are also regulated on the post-transcriptional level and thus show the limitation of using gene expression as a proxy for the abundance of the active form of TFs.

**Figure 5.**
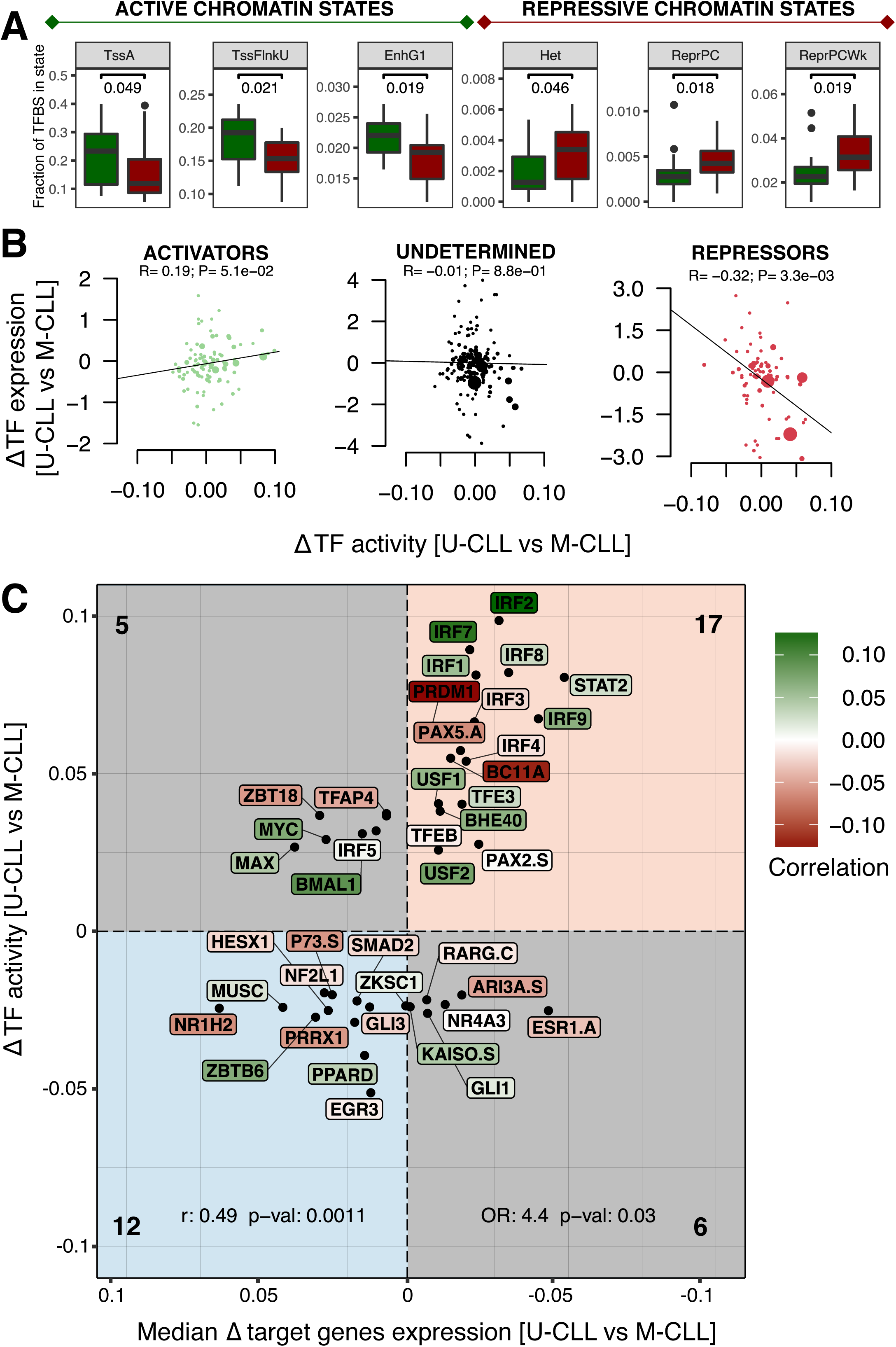
Validations for the activator and repressor classification and downstream analyses. **(a)** Boxplots showing the fraction TFBS overlapping with specific chromatin states as defined by *chromHMM* (Roadmap Epigenomics Consortium et al., 2015) for all activators (green) and repressors (red) are shown. Only *chromHMM* states with significant differences (Wilcoxon test; p-value < 0.05) between activators and repressors are displayed. **(b)** Pearson correlation of the log2 fold changes from RNA-Seq and differential TF activity for activators (green, left), undetermined TFs (black, middle), and repressors (red, right). Only expressed TFs are shown. **(c)** Correlation of differential TF activity and median of the differential target gene expression of U-CLL against M-CLL. The x-axis shows the median target gene (TG) log2 fold-change, the y-axis denotes the differential TF activity. Each TF label is colored based on its activator/repressor status (green/red) on a continuous scale (dark to light) based on the correlation strength (see Fig. 4d). (OR=odds ratio, r=Pearson correlation coefficient).

### Case study II: Applying diffTF to small scale multiomics dataset

To assess the applicability of diffTF to small datasets, we decided to apply it to the well-studied mouse hematopoietic system. We generated ATAC-Seq and RNA-Seq profiles of multipotent progenitor cells (MPP; Lin^-^cKit^+^Sca1^+^; CD150^-^CD48^+^), an early hematopoietic progenitor population capable of supporting multilineage blood production (Sun et al., 2014), as well as the more differentiated and myeloid-restricted granulocyte-monocyte progenitors (GMP; Lin^-^cKit^+^Sca1^-^; CD16/32^+^). The profiles obtained were processed using an in-house ATAC-Seq pipeline and *diffTF* (using the analytical procedure due to the small number of samples) to identify TFs that are differentially active between MPP and GMP cells (see Online Methods). Due to the large number of significant TFs, reflecting the high diversity between the two cell types, we used RNA-Seq data to filter out non-expressed TFs. The differential signal is dominated by an increase activity of the members of the well-known class of master regulators of myelopoiesis, the CEBP family (C/ebpα,-β,-δ,-ε,-γ) in GMPs (**Fig. 6a**, **Suppl. Fig. 16**). In addition, we observed higher activity of the MYC/MYB factors, which are known to be exclusively active in the GMPs (Baker et al., 2014) and in NFIL3, which is involved in the generation of natural killer cells (Gascoyne et al., 2009). Conversely, MPPs show a higher activity for IRF/STAT, ZEB1 and ITF2 (part of the Wnt signaling) as well as TFs from the Homeodomain (HXB7,HXA10) and Forkhead (FOXO3) families, all of which are associated with self-renewal of hematopoietic stem cells (Sands et al., 2013).

**Figure 6.**
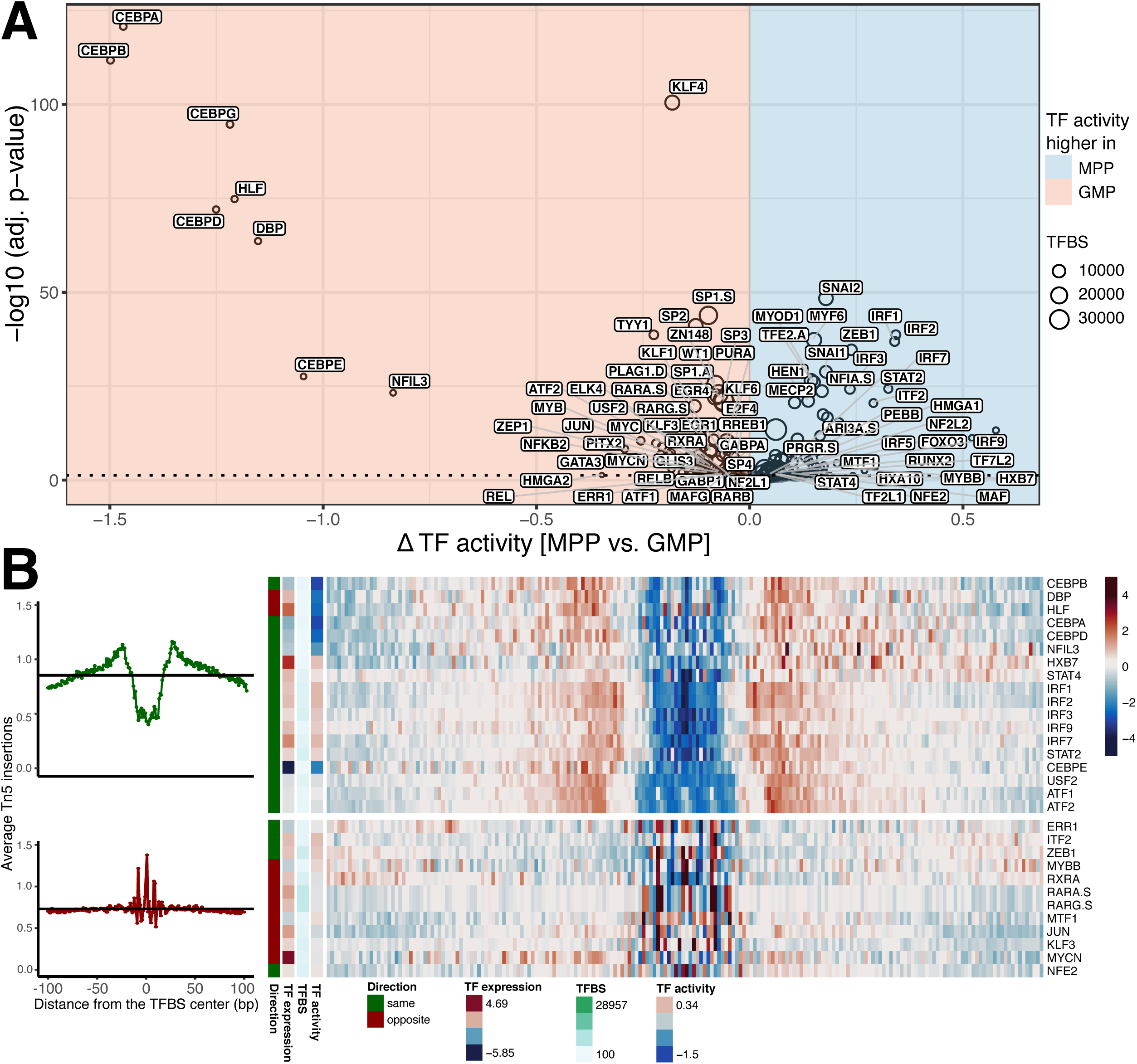
diffTF recapitulates known TFs that drive the differentiation from MPP to GMP and shows a similar activator/repressor cluster as in the CLL data. **(a)** Volcano plot of differential TF activity between MPP (n=4) and GMP (n=4) cells. Due to the high number of significant TFs only the most significant are labeled. The full list is available in Suppl. Table 4. **(b)** The footprints are for TFs in two selected clusters that represent the activators and repressors, respectively are shown as heatmap (right) and aggregate plots (left; see **Suppl. Fig. 17** for the full heatmap). Only TFs that were significantly differentially active and all significantly differentially expressed (adj. p-value < 0.05 for both) are displayed. Colors represent footprint strength, while white denotes the value of the genomic background in the consensus peakset. Clusters were defined using hierarchical clustering with the *ward.D2* method (clustering tree omitted for clarity)(see **Suppl. Fig. 17**). For the cluster summary footprints at the left, we divided each footprint value by the mean value of each cluster to highlight the differences in the surrounding chromatin structure. The direction of TF expression and TF activity is in analogy to what is described in the text. “Direction” denotes whether expression and TF activity have the same or opposite sign.

The small number of samples made the correlation-based classification of the TFs into activators and repressors unreliable. Therefore, we devised a second - less quantitative - approach for activator/repressor classification that is based on the TF footprint and differential RNA-Seq expression. In short, we determined whether TF activity and expression level co-vary in the same direction and combined this with visual inspection of the pattern of its footprint (**Suppl. Fig. 17**). This allowed us to identify the set of activators that showed a clear activating footprint as observed for the Class I TFs in the CLL data (**Fig. 6b**). Similar to the Class II and III for CLL, the pattern was less clear for the repressor footprint clusters, which contain both potential repressors and activators. Interestingly, the most differentially active TFs between MPP and GMP are mainly classified as activators (CEBPs, NFIL3, IRFs) or have mixed evidence (i.e. activator footprint, but inconsistent directionality of expression and activity such as DBP and HLF). The most differentially active repressor we identified is JUN, whose difference in activity is far below the activators, indicating that the differentiation process from MPP to GMP is mainly driven by transcriptional activators.

Overall, these results show that *diffTF* is able to identify the known TFs that drive the differentiation from MPP to GMP, thus demonstrating its power to detect signals also for a small number of samples. Furthermore, we show how a qualitative classification scheme of TFs into activators and repressors that is primarily based on TF footprints can be useful in comparisons where the small number of samples does not allow a correlation-based classification.

### Comparison with similar tools

We compared the few tools with a similar focus (Baek et al., 2017; Heinz et al., 2010; Schep et al., 2017) with *diffTF* (**Suppl. Table 5**) both qualitatively (*chromVAR, BagFooT, HOMER*) and quantitatively (*HOMER* and *chromVAR*). Overall, *diffTF* provides a more flexible and tailored analysis framework due to the extensive choice of parameters, diagnostic plots, TFBS-specific results, visualizations, and pipeline adjustability. As mentioned above, it is unique in its ability to directly integrate RNA-Seq with ATAC-Seq data to classify TFs into activators and repressors. Due to its flexibility, *diffTF* is computationally expensive, and we provide detailed instructions on memory and time requirements in the documentation.

We first compared *diffTF* with a more traditional TF motif analysis such as *HOMER* (Heinz et al., *2010)*, which looks at motif enrichment in a set of differentially accessible peaks. Strikingly, no enriched motifs were found in M-CLL with *HOMER*, while the few discovered in U-CLL correlated significantly with differential TF activity as computed by *diffTF* (**Suppl. Fig. 18,** see Methods). This analysis highlights the power of *diffTF* to capture more signal than standard motif enrichment approaches.

To compare *diffTF* with an approach that is also based on TF activities we chose *chromVAR* and *BaGFoot* (Baek et al., 2017). We were unfortunately unable to run and adopt the BagFoot workflow for our CLL data due to missing example files, an incomplete documentation and unresponsiveness from the authors. *For chromVAR*, the results correlate very well overall, with correlation coefficients between 0.75 and 0.93 (Pearson) and 0.69-0.88 (Spearman), depending on the set of TFs (i.e., all TFs or only those deemed significant by *diffTF*, therefore predominantly removing TFs with low signal) and whether *chromVAR* deviations or deviation scores are compared against (**Suppl. Fig. 19a-b**, see Methods). Differences likely arise due to distinct methodological divergences such as comparing fold-changes for peaks (*chromVAR*) vs. binding sites (*diffTF*) or whether to compare the TF-specific effect against the mean effect across all TFs (*diffTF*) or not (*chromVAR*; see also Methods and **Suppl. Fig. 19c-f**). However, *diffTF* goes one step beyond the currently available methods by classifying TFs based on their mode of regulation - activator or repressor, thus providing important additional insights into their molecular function.

## DISCUSSION

We presented a genome-wide method for quantifying differences in differential TF activity for a large set of TFs simultaneously, and for classifying them into their molecular mode of action as transcriptional activators or repressors. The method is available for download at https://git.embl.de/grp-zaugg/diffTF along with a comprehensive documentation and example data.

We have shown in two case studies that *diffTF* is able to recover a change in activity for the TFs expected to drive the biological processes, thus demonstrating the biological validity of the method. In addition, we have extensively tested and demonstrated the technical robustness of *diffTF*. In particular, we have shown that *diffTF* is able to overcome the inherent noisiness of TF binding site predictions by aggregating data across all putative binding sites.

Calculating differential TF activity based on aggregating signal across the genome has been proposed before based on the expression of putative target genes of a certain TF (Boorsma et al., 2008; Bussemaker et al., 2001). Using the effect on chromatin instead of expression has several advantages: first chromatin is a much simpler trait since gene expression is the sum of transcription and degradation, thus increasing the power to detect differences. Second, there are much more peaks than genes, thus allowing for better statistics and signal to noise ratio. Finally, the effect on chromatin is much more local than on gene expression – in particular in mammalian genomes, where genes can be regulated by distal enhancers. We have compared differential TF activity calculated based on the average expression change of the target genes to the output of *diffTF* and found that while the direction of activity between both methods is highly correlated, the signal is much lower when using the expression of target genes instead of chromatin accessibility at putative binding sites.

To demonstrate the power of *diffTF* for large but heterogeneous datasets, we have applied it to identify and characterise differences between M-CLL and U-CLL from a publicly available dataset of ATAC-Seq (1bn reads, 52 patients) and RNA-Seq. It is noteworthy that a TF motif enrichment analysis on the significantly differentially accessible peaks did not reveal any factor to be significantly enriched in M-CLL, indicating that in this case (as probably in many patient-control studies) the key TFs are not necessarily switching their target enhancers and promoters on and off, but rather mis-regulating many regions to a lesser extent. The advantage of *diffTF* is that it can detect a slight shift in activity of the TF even if the signal differences at each binding site a very low and rarely significant. It does so by averaging across all of a TFs putative binding sites and is therefore more powerful than conventional enrichment analyses.

We have devised an approach within the *diffTF* framework to classify TFs into activators and repressors based on the correlation of their expression level (RNA-Seq) and the activity of their putative binding sites (ATAC-Seq). This information is highly relevant when interpreting the action of TFs since it is important to know whether an upregulation of a TF would have a positive or negative effect on chromatin (and therefore transcriptional) activity. Notably, this classification could work even for datasets for which insufficient RNA-Seq data are available – as we have shown for the MPP-GMP case – by jointly investigating TF footprints, differential expression of the TF and differential TF activity.

It is important to note that TFs can often act as activators and repressors at different genomic loci e.g. depending on their cofactors, whereas here we predict their main mode of action based on the mean effect across all their predicted binding sites, and thereby lose any information about bifunctionality. Furthermore, since the classification is based on correlations, it is heavily dependent on variation in the RNA-Seq signal across individuals, thus biasing the TFs that can be classified towards those that are variable across individuals. As a consequence, TFs whose post-transcriptional regulation is not reflected in their transcript abundance will not get classified correctly. Another potential misclassification may happen because of the similarity of PWMs within a cluster, which makes it difficult to distinguish the exact effect of one TF while its expression level is uniquely defined. As an example we cite PRDM1, which is part of IRF family (cluster 40) and classified as very strong repressor, its footprint however looks more similar to a typical activator (data not shown), suggesting that it is not PRDM1 driving the ATAC-Seq signal in this case, but the IRFs. Thus, for distinguishing the functional roles of TFs from the same cluster/family further biochemical experiments will be needed. Despite these potential pitfalls, *diffTF* provides unique insights into the molecular mechanism of TFs on a global level.

Since many ATAC-seq experiments have a rather low number of samples, we also assessed the power of *diffTF* to uncover biology in small (but more controlled) experiments. In particular, we have performed a *diffTF* analysis to compare murine MPP and GMP (4 replicates each). Again we identified the major TFs driving the differentiation, and were able to qualitatively classify TFs into activators and repressors - in a correlation-independent approach. This classification revealed that the bulk of the change in chromatin accessibility during the differentiation from MPP to GMP is driven by activators. This case-study demonstrates the applicability of *diffTF* to small-scale data.

While similar methods have been proposed for analysing ATAC-Seq data (Baek et al., 2017; Schep et al., 2017), our method has several advantages when dealing with bulk ATAC-Seq data and can also be used for histone mark ChIP-Seq data: (i) Unlike other methods that calculate the background theoretically based on the genome-wide read depth, *diffTF* is insensitive to sequence and locus dependent biases since we calculate a fold-change between conditions for each region, thus normalizing for local read depth biases. This is particularly advantageous for detecting small changes such as between two heterogeneous cohorts in patient-control studies. (ii) *diffTF* allows integration with matching RNA-Seq data to classify TFs into activators and repressors in a fully data-driven approach within the same analysis framework. Such classifications are a significant help when interpreting the effects of up/down regulation of a particular factors. (iii) *diffTF* provides the fold-change value for each TFBS which allows for easy retrospective follow-up analysis, e.g. identifying the most differential regions regulated by a specific set of TFs. (iv) Finally, our method might allow to analyse time-course data in an additive manner by calculating the overall change of slope for each TF (see Methods).

Overall, with *diffTF* we present a multiomics data integration strategy of ATAC-Seq and RNA-Seq data that calculates differential TF activity across conditions and classifies TFs based on their molecular mode of action into activators and repressors. With this, *diffTF* can aid in formulating testable hypotheses and ultimately improve the understanding of regulatory mechanisms that are driving the differences in cell state on a systems-wide scale.

## METHODS

Methods, including statements of data availability and any associated accession codes and references, are available in the online version of the paper.

## Supporting information

## ACKNOWLEDGMENTS

We thank Bernd Klaus for help with the statistical part and EMBL for funding. A.R.P is recipient of a postdoctoral fellowship granted by Fundación Ramón Areces. G.P. was supported by the Otto Bayer Scholarship from Bayer Foundations.

## AUTHOR CONTRIBUTIONS

J.Z. and I.B. conceived the study, C.A. and I.B. developed the computational framework and performed the analyses, A.R.P. and G.P. contributed to the development and analysis, K.D.R. performed the experiments, K.H. supervised the experiments, J.Z. supervised the study and C.A., I.B., and J.Z. wrote the manuscript.

