## Supplementary material for "Quantification of differential transcription factor activity and multiomics-based classification into activators and repressors: *diffTF*"

### ONLINE METHODS

**Data used in analyses**

**Tools used**

We used the workflow manager *Snakemake (Köster and Rahmann, 2012)* 5.0 for all described pipelines as well as the *Bioconda* project (Grüning et al., 2018) to maintain and handle software via the *Conda* package manager (https://conda.io). The ATAC-Seq preprocessing pipeline uses *bedtools* 2.26.0 (Quinlan and Hall, 2010), *samtools* 1.4, *Bowtie2* 2.3.0 (Langmead and Salzberg, 2012), *deepTools* 2.5.0 (Ramírez et al., 2014), *FastQC* 0.11.5 (Andrews, 2010), *GATK* 3.7(McKenna et al., 2010), *MACS2* 2.1.1 (Zhang et al., 2008), *multiqc* 0.9 (Ewels et al., 2016), *Subread* 1.6.0 (Liao et al., 2013), *Picard* tools 2.9.0, *Trimmomatic* 0.36 (Bolger et al., 2014), *PWMScan* (Ambrosini et al., 2018) web interface (http://ccg.vital-it.ch/pwmtools) and standalone release 1.1.1 (https://sourceforge.net/projects/pwmscan) and R version 3.4. In addition, a number of R/Bioconductor (Huber et al., 2015) packages were used, the most important of which are described below. For a full list of used packages and their corresponding versions, see the Github repository at https://git.embl.de/grp-zaugg/diffTF. For the various additional analyses that we performed, we used the following tools: *HOMER* v4.9 (Heinz et al., 2010), *BiasAway* 0.96 (Worsley Hunt et al., 2014), *MEME* suite 4.11 (Bailey et al., 2009) and the R/Bioconductor *chromVAR* package (Schep et al., 2017).

**ATAC-Seq processing**

The *diffTF* pipeline requires properly processed and quality-controlled ATAC-Seq data. To obtain these, we developed an in-house *Snakemake* pipeline. The pipeline starts with raw fastq files and integrates multiple steps for quality control, adapter trimming, alignment, as well as general and ATAC-Seq specific post-alignment filtering and processing steps.

In the following we describe the pipeline using the default parameters. *FastQC* is used to assess the sequence quality. Foreign sequences from the Nextera Transposase agent are removed with *Trimmomatic*, using the parameters *ILLUMINACLIP:NexteraPE-PE.fa:1:30:4:1:true TRAILING:3 MINLEN:10*. Alignment is performed with *Bowtie2* with *-X 2000* (maximal fragment length), *--very-sensitive* and against hg19, followed by various cleaning steps (*Picard* tools *CleanSam, FixMateInformation*, *AddOrReplaceReadGroups*, and *ReorderSam*) and base quality recalibration using GATK with known variants taken from the provided *GATK* bundle for hg19 (SNPs: *dbSNP* version 138, Indels: *Mills_and_1000G_gold_standard.indels*). This allows us to detect and correct systematic errors made by the sequencer when estimating the quality score of each base call, thereby increasing data quality.

The pipeline then performs various cleaning and filtering steps: (1) removing mitochondrial reads and reads from non-assembled contigs or alternative haplotypes, (2) filtering reads with a mapping quality below a user-specified threshold (default = 10), (3) marking and removing duplicate reads with *Picard* tools, (4) adjusting read start sites as described previously(Buenrostro et al., 2013) (4 bp on the forward and 5 bp on the reverse strand) and (5) removing reads with insertions or deletions using *samtools* (Suppl. Fig. 6a-b).

Lastly, a GC bias diagnosis and correction using *deepTools* and Benjamini's method (Benjamini and Speed, 2012) is run for each sample. This helps to assess the severity of the GC bias in the data, namely that of DNA polymerases used for PCR-based amplifications during the library preparation, which usually results in artificially higher read counts for GC rich regions (Suppl. Fig. 6d-e).

The output of the pipeline can be obtained for the (1) original and (2) GC corrected data as well as on the level of (3) individual files (i.e., one file per replicate) or (4) pooled replicates (i.e., one file per sample). Downstream analyses for these four classes of files involve peak calling using MACS2 with user-adjustable stringencies and parameters and removal of blacklisted regions. Finally the pipeline outputs summary statistics and additional files and plots (coverage files for visualization, transcription start site enrichment, sample-specific fragment length distributions (Suppl. Fig. 6c), library complexity measures, PCA (Suppl. Fig. 7), sample correlations).

#### Description of the *diffTF* analysis workflow

In summary, *diffTF* calculates a differential "activity" between two or more conditions for each TF by comparing the distribution of fold-change differences across all binding sites of a TF to all binding sites from all other TFs. The algorithm is split into 7 steps as described in the following:

*1. Generating a consensus peak set*

Our approach is based on analyzing read counts in peaks, which requires a consensus peak set across input samples. This can either be user-provided or it is generated from the sample-specific peaks. For the latter, consensus peaks are generated with the function *dba.peakset* in the *DiffBind* (Ross-Innes et al., 2012) R/Bioconductor package (using the parameter *minOverlap* to define the number of samples within which a peak should be present). We then retain only peaks from genuine autosomes, thereby filtering sex chromosomes, non-assembled contigs as well as alternative haplotypes. The consensus peak set is finally sorted by coordinate to speed up subsequent computations.

*2. Scanning of TF binding sites*

For each TF of interest, *diffTF* needs a set of TF binding sites (TFBS). To generate them, we used the *HOCOMOCO* database, which provides TF binding models (PWMs) that are collected from numerous ChIP-Seq experiments for 640 TFs in human and 423 in mouse (Supplementary Table 3). We used these PWMs to scan the hg19 genome using the *PWMscan* web interface and the standalone version to obtain all putative binding sites for each TF (cutoff p-value - 0.00001, background base composition - 0.29;0.21;0.21;0.29). We finally sort the TFBS for each TF by coordinates. For user convenience, we provide this sorted list for both human and mouse in the Git repository. However, *diffTF* is not limited to any specific database and any tool can be used to predict TFBS.

*3. Differential analysis for the consensus peakset*

To calculate the fold-change between the two conditions across each peak, we first obtain the counts for the consensus peakset for each sample using *featureCounts* from the *Subread* package with the options -p -B -d 0 -D 2000 -C -Q 10 -O -s 0*.* We then employ *DESeq2* for count normalization, using a cyclic loess approach as implemented in the normOffsets function of *csaw* (Lun and Smyth, 2016) from R/Bioconductor to alleviate potential systematic biases between the samples. Importantly, unrestricted design formulas can be incorporated into *diffTF*, making it therefore a very flexible approach if additional co-variariates should be incorporated to increase statistical power. In addition to the two conditions comparison as described above (e.g., mutated vs. unmutated, wildtype vs. mutant, young vs. old), *diffTF* can also incorporate design formulas for which the predictor variable is continuously-valued rather than binary to analyze time-course data (e.g., age, differentiation time, etc). The reported log2 fold change is then per unit of change of that variable. That is, TFs with a negative differential TF activity have a negative slope per unit of change of that variable, while TFs with a positive TF activity have a positive one. To check the normalization results, we generate four types of diagnostic plots: (1) MA plots, (2) density plots of normalized and non-normalized counts, (3) mean-average plots (average of the log–transformed counts vs the fold-change per peak) for each of the sample pairs and (4) mean SD plots (row standard deviations versus row means). See Suppl. Fig. 5a-b for an example.

*4. Signal extraction for each TFBS*

We then filter for TFBS that overlap with the consensus peak set only while allowing for multiple TFBS per peak (using the program *bedtools intersect* with options *-wa -wb*). Each TFBS is then extended by 100 bp (user-adjustable) in both directions (motif extension), followed by extraction of read counts for each sample (*featureCounts*, see above for the parameters used).

*5. Calculation of accessibility fold-change for each TFBS*

To avoid biases and dependencies based on TFBS clustering within peaks, we then select the TFBS per TF per peak with the highest average read count across all samples. On this set we perform a differential “accessibility” analysis using a standard *limma* workflow employing the functions *lmFit* and *eBayes* independently for each TF for the same design formula as used in step 3 (that is, either a model contrasting two conditions or a continuously-valued model). For count normalization, we use the consensus peak-derived normalization factors from step 3. As a result we obtain a log2 fold-change value for each selected TFBS per TF per peak, and in addition we provide various diagnostic plots for each TF (same as in step 3, with additional ECDF and density plots of the log2FC values, Suppl. Fig. 5c-d).

*6. GC binning and calculation of differential TF activity values*

(a) GC binning

To reduce biological biases based on differential effects depending on GC content of the local environment of a TF, we first obtain the GC content of each extended TFBS with *bedtools nuc.* We then group TFBS into 10 bins (user-adjustable) based on the GC content of their extended TFBS (e.g., +/-100bp).

(b) Estimation of the differential TF activity

For each bin containing more than 20 TFBS for a given TF, we compare the TF-specific distribution of log2 fold-changes against a background of log2 fold-changes of all TFBS from all other TFs of the same GC bin. We then define the difference in means as mean difference for each bin. Finally, for each TF, the differential TF activity is calculated as the weighted arithmetic mean across all mean difference values for the bins with sufficient data, weighing each value from each bin by the fraction of TFBS it contains in that TF such that all weights sum to 1.

*7. Estimation of significance for differential activity for each TF*

The significance and magnitude of the TF-specific TF activity values can be difficult to interpret. To assess their statistical significance, we employ a permutation approach to derive empirical p-values (hereafter called the empirical approach). We rerun steps 3-6 for a total of 1000 times (user-adjustable) with permuted input data (permutation of condition labels) and then calculate an empirical two-sided p-value per TF by comparing the real value with the distribution from the permutations and calculating the proportion of sampled permutations for which the absolute differential TF activity is larger (Suppl. Fig. 4a-c). To validate our procedure, we also plot the density of the TF activity across all permutations and compare it to the real data, which expectedly gives a distribution that is tightly centered around 0 for the permuted data while the real data show a shift as well as heavier tails (Suppl. Fig. 4d). We finally perform multiple testing correction using Benjamini-Hochberg (Benjamini et al., 1995).

If the number of samples is small or only very few samples come from one condition, the number of possible permutations is also small, therefore making it difficult for the permutation-based approach to accurately assess significance. In such a case, we offer a modified version of the described pipeline that we call the analytical approach: First, we use *DESeq2* instead of *limma* to calculate fold-changes throughout the pipeline for the TF-specific steps, which is slower but better suited for a small number of samples. Second, instead of employing permutations, we run a modified procedure from step 7 onwards to assess the statistical significance of the differential TF activity: To reduce the dependency of the p-value on the sample size (since the number of TFBS can range between a few dozen and multiple tens of thousands depending on the TF), we first perform a Welch Two Sample t-test for each bin and calculate the overall significance by converting the resulting T-scores to z-scores, which allows to summarize them across the bins and convert them to one p-value per TF (Suppl. Fig. 4f-g). Because we calculate the weighted mean out of the individual T-scores and therefore end up having a weighted sampling distribution of the mean, we however need to estimate its expected variance.

(a) Estimation of the variance

To estimate the variance of the weighted mean T-scores for each TF we use the following formula (see https://en.wikipedia.org/wiki/Propagation_of_uncertainty#Linear_combinations) in which $w_i$ and $x_i$ represent the weight and T-statistic value from the t-test of a particular bin *i*, $Cov(x_i, x_j)$ the covariance of two particular bins *i* and *j*, and $var(x_i)$ the variance of the T-statistic value for $x_i$ :

$$var(\underline{x}) = \sum_{i=1}^{n} var\left(\frac{w_i}{\sum_{k=1}^{n} w_k} x_i\right) + 2 \sum\sum_{1 \le i < j \le n} w_i\, w_j\, Cov(x_i, x_j)$$ (weights sum up to 1)

$$= \sum_{i=1}^{n} var(w_i * x_i) + 2 \sum\sum_{1 \le i < j \le n} w_i\, w_j\, Cov(x_i, x_j)$$ (if all values are scaled by a constant for the variance, it is scaled by its square)

$$= \sum_{i=1}^{n} w_i^{\,2}\, var(x_i) + 2 \sum\sum_{1 \le i < j \le n} w_i\, w_j\, Cov(x_i, x_j)$$

This is done to prevent a systematic misestimation of the variance of the $x_i$ terms by assuming it to be 1 for all bins. To properly estimate $var(x)$, we employ a bootstrap approach using the *boot* library in R with a user-adjustable number of bootstrap replicates (default 10,000), with resampling the bin-specific data and then performing the t-test against the full sample as described above. We then calculate the variance $var(x_i)$ of the bootstrapped T-scores for each bin. Since the T-scores across the bins are not necessarily independent, we also correct their variance using their pairwise covariance, which is is estimated with the term $Cov(x_i, x_j)$ using the bootstraps values for each bin. Across different analyses and datasets, we generally observed that covariances are predominantly negligible, but for some TFs and bins, they differ significantly from 0 and therefore it is important to incorporate them. A detailed analysis of the covariances $Cov(x_i, x_j)$ between pairs of bins of the TF-specific distribution of log2 fold-changes against a background of log2 fold-changes from a randomly selected set of TFBS of the same GC bin (see step 6 and the formula in step 7a) revealed that covariances are negligible in almost all cases. From the 9,099 covariances across all TF and pairs of bins, only 19 cases exceeded an absolute value of 0.05 and only one 0.2, while over 91% had absolute values of smaller than 0.02 (data not shown). We verified the validity of this procedure with a simulation script (available upon request) that shows that p-values are well behaved for random data (Suppl. Fig. 4i).

*(b) Calculation of one p-value per TF*

To obtain a p-value for each TF, we centralize the distribution of the weighted T scores across all TF by subtracting its mean, which we obtain from the maximum likelihood estimate of the distribution mean using the locfdr function from the locfdr package in R. We then calculate p-values out of z-scores based on the TF-specific variance calculated above. We finally perform multiple testing correction using Benjamini-Hochberg method (Benjamini et al., 1995).

**Validation of the analytical approach**

We used the GMP/MPP data to verify the validity of our analytical approach by comparing the real signal from the GMP vs. MPP comparison with an array of additional analyses: We first run *diffTF* within GMP and MPP, respectively, and repeated this for all possible pairwise combinations (2 vs. 2). In addition, we run two types of controls: One regular *diffTF* analyses of GMP vs. MPP in which we shuffled the TFBS before the binning and calculation of statistical significance and one in-silico simulation based on random data that mimics precisely the *diffTF* workflow while incorporating some of the GMP vs MPP specific parameters (see Methods). Overall, we found that comparing GMP vs MPP gives the strongest signal, while comparing only within GMP or MPP cells still yields signal albeit on a much smaller scale, while the two types of control indeed showed no signal (Suppl. Fig. 4i)**.** The remaining signal within cell types is a tribute to the high sensitivity of *diffTF* to pick up small signal and reflects the inter-individual differences between biological replicates. We therefore recommend for the analytical approach to perform at least one within-sample analysis to obtain an idea of the expected signal based on inter-individual variation of biological replicates.

**Guidance on the number of permutations and whether to sue the analytical or permutation-based approach**

As explained before, *diffTF* offers two procedures to calculate significance , and we want to give some guidance what the significance in each procedure means, when to use which, and how to adjust the parameters.

(1) Permutation-based approach

In this approach, the resulting significance value captures the significance of the effect size (that is, the TF activity) for the real data as compared to permuted one. Importantly, this significance measure appears independent of the number of binding sites per TF (r=0.026, Spearman, see also Suppl. Fig. 4e), and should be used when the number of samples is large enough to perform a reasonable number of permutations. The range of p-values will be determined by the number of permutations (smallest p-value equals to 1/{# of permutations}) and we recommend at least 1000 permutations (as we have done for the CLL dataset). The number of possible permutations can be calculated with the binomial coefficient $\binom{n}{k}$, with *n* being the total number of samples across the two conditions and *k* either of the two total number of samples for one of the two conditions. For example, when comparing 5 GMP against 4 MPP samples, there are $\binom{9}{5}$ = $\binom{9}{4}$ = 126 possible permutations. The minimum raw p-value $p_{min}$ > 0 is then $\frac{1}{126} \sim$ 0.008. It is computationally more expensive than the analytical approach.

(2) Analytical approach

Here, the resulting significance value is calculated by performing t-tests for each TF and GC bin (see methods above). Given that both the p-value and to a lesser extend also the

T statistic depend on the sample size (i.e. the number of binding sites *n*), we checked whether this dependence would explain much of the signal. However, for both CLL and GMP/MPP, we found highly significant TFs with lower number of binding sites despite the correlations of p-value and number binding sites (r= -0.158 and r= -0.409, respectively, Spearman), thus showing that the biological signal captured by *diffTF* surpasses the theoretical bias (Suppl. Fig. 4h).

If the number of samples is too small to allow a reasonable number of permutations in the permutation-based approach or computation time is important, this approach can be used. The number of bootstraps for the variance estimation of the T scores should be set to a high value, and we recommend values of at least 1,000 or even 10,000. Computation time, however, increases with the number of bootstraps. For the CLL data, the estimated bootstrap variance approaches the expected value 1 very quickly even for bins with low numbers of TFBS (the minimum is set to 20).

**CLL ATAC-Seq processing**

We run the ATAC-Seq pipeline on the 52 CLL samples as described above with the indicated default parameter values. The distribution of the number of the fraction of retained and total number of reads across all samples throughout the course of the pipeline showed the typical pattern, namely that reads were filtered mainly because they were of either of mitochondrial origin or duplicates (Suppl. Fig. 6a-b). The fragment length distributions also showed the typical and expected pattern, with peaks at expected lengths (e.g, mono- and dinucleotides) (Suppl. Fig. 5c). Lastly, they showed clearly that read counts were driven by the GC content (see Supplementary Figure 6d for a randomly chosen example), which we eliminated using the GC correction as described. The diagnostic plots after the GC correction then expectedly indicated no GC dependency (see Suppl. Fig. 6e). We also performed a PCA analysis of the consensus peak regions (Suppl. Fig. 7).

**Running *diffTF* on the CLL data**

We used 52 samples out of 88 available ATAC-Seq samples from the chronic lymphocytic leukemia data. We excluded 4 samples because there mutational status was unknown. Out of 84 remaining samples, we used only one replicate per sample ("_1"), which resulted in a total of 52 samples (25 U-CLL and 27 M-CLL, see also Supplementary Table 3) in order to not bias the analysis by overrepresenting particular individuals due to the varying number of replicates per sample. As discussed above we used the GC-corrected BAM files hereafter. A PCA analysis of these peak regions showed a clear separation of the samples based on their IgHV mutation status, but not on any other potentially confounding factor like batch, IGVH homology, gender, the patient age at data collection or the patient age when diagnosed (Suppl. Fig. 7). We generated a consensus peakset as described above using 5 samples as the minimum overlap. This retained 48,065 consensus peaks which were used subsequently. As motif extension, we added 100 bp on each side of the predicted *in silico* TFBS. As parameters for the peaks and TF-specific differential analysis, we used the design formula: *"~ batch* + *mutationStatus*" and *fitType* = *'local'*. Here, *batch* and *mutationStatus* are metadata that are available for each sample and that refer to sample batch and its mutation status (either mutated or unmutated), respectively. Based on the discussion in step 6, we decided to use 10 bins for our GC binning approach (that is, ranging from 0-10% GC content up to 90-100%).

**Comparison of *diffTF* results based on predicted vs. ChIP-Seq validated TFBS**

We used TF binding data gathered by *ReMap* from human ChIP-seq experiments and intersected them with *in silico* predicted TFBS using TF binding models from the HOCOMOCO database. We then split the 157 common TFs by whether or not they overlapped with the *ReMap* CHIP-seq data and rerun the *diffTF* pipeline. Finally, we correlated these two groups with and without ChIP-seq validation using Pearson correlation. The correlation between the TFBS that were present vs. not present in *ReMap* was 0.54 (P=3.5e-10) (Suppl. Fig. 8).

**Assessing impact of TF motif scanning parameters on *diffTF* results**

For the purpose of assessing the effect of the genome-wide TFBS scanning on *diffTF*, we varied the p-value cutoff in *PWMScan* (0.00005, 0.00001, 0.000001) using the default background base composition of 0.29;0.21;0.21;0.29 as well as varying the background base composition (0.27;0.23;0.23;0.27) (Suppl. Fig. 9). The former resembles the human genome nucleosome composition, the latter the base composition from the ATAC-Seq peak regions only. For this, we generated the FASTA file of the consensus peak set with `*bedtools getfasta*` and then calculated the background nucleotide composition with `*fasta-get-markov*` from the *MEME* suite. PWMs from *HOCOMOCO* and PFMs from *JASPAR* were converted into integer log likelihoods using `*pwm_convert*` (setting -f as "real" and "jaspar" for PWM and PFM, respectively). Scanning was performed using the script `*pwm_bowtie_wrapper*`. See Suppl. Fig. 9.

**_diffTF_ robustness analysis with respect to internal parameters for the CLL data**

As discussed above, we verified the robustness against internal parameters such as the number of permutations or the number of bootstraps (for the analytical significance calculations) and found that *diffTF* results are not dependent on these. Most importantly, we also systematically varied the default motif extension size from the default 100 bp to 0, 50, 200, 400, and 600 bp to investigate the effect of the extended TFBS on the *diffTF* results (Suppl. Fig. 10). In summary, the TF activity differences were very similar, with correlations above 0.9 and 0.86 for motif extension sizes from 50-600 for Pearson and Spearman, respectively. Only an extension size of 0 showed weaker correlation, albeit the values were also surpassing 0.76 and 0.71, respectively. For almost all TFs, the TF activity difference estimates were consistent across motif extension sizes, with only a few TFs changing directions from positive to negative values or vice versa. For large extension sizes, the magnitude of the differential TF activity.decreased overall. Running *diffTF* with no extension size or too large values (>500), therefore, seems inappropriate as it either excludes too much signal directly adjacent to the predicted TFBS or the resulting extended TFBS are becoming too wide.

Raw and adjusted p-values had more differences when varying the extension size, with correlations above 0.53 and 0.58 for extension sizes from 50-600 for Pearson and Spearman, respectively. Not using any motif extension showed again weaker correlation with values at least being 0.28 and 0.35, respectively. In conclusion, the raw and adjusted p-value comparison across extension sizes identified mainly three clusters of TF: a small number of TFs gaining or losing significance with increasing extension size, respectively, and the majority of TFs being invariant for it. 67% of all TFs that we identified as being significant remained significant throughout the majority across all extension sizes, while almost 95% were significant for at least one other extension size.

**Assessing dependence of _diffTF_ results on differential signal strength for CLL data**

In order to assess the effect of differentially accessible (DA) peaks on *diffTF*, we generated a peak set excluding DA peaks. For this, we used *DiffBind* (with design ~ *batch_number* + *Condition*) and identified 389 and 3569 DA peaks for the mutated and unmutated condition, respectively, which we then excluded from the original peak set. We then run *diffTF* with standard parameters (described above) and correlated the results (see Fig. 2f).

**Assessment of the power of *diffTF* with respect to sample size and sequencing depth for CLL data**

To test the power of our approach and the dependence on the (i) read depth (coverage) of the BAM files and (ii) number of samples for each of the two conditions, we performed a subsampling procedure in which we varied both read depth and sample size. We used the full 84 sample CLL dataset for this in order to maximize the number of samples. To vary the read depth, we used the original, non-GC corrected CLL data that is produced as part of the output of our ATAC-Seq pipeline and then randomly generated downsampled BAM files using Picard *DownsampleSam* with random seeds and PROBABILITY=k, with k ranging from 0.75, 0.50, 0.25, 0.125, 0.06, 0.02 to 0.01. For each of these fractions, we (1) determined the median number of reads these fractions correspond to across all samples, thereby giving an estimate of the required number of processed reads to produce a particular accuracy and (2) run the *diffTF* pipeline with the addition of also varying the number of samples per condition while maintaining the original ratio of the two conditions (around 60 % to 40%). The latter we varied from the full data 34+50 down to 30+45, 25+37, 20+30, 15+23, 10+15, and down from 9+14 in steps of 1 to 3+5, with the first and second number denoting the number of distinct samples with the condition unmutated and mutated, respectively. For each of these cases, we in addition performed 50 repetitions to minimize sampling noise. We then compared the values from the full dataset with each of the two-way subsampled data and evaluated the results using the following measures: (1) the fraction of TFs that show the same direction of change (that is, either positive or negative) for the differential TF activity as in the full data, and (2) the correlation of the differential TF activity with respect to the full data. For (1), we split all TFs into three equally sized bins using the 33% and 66% quantile threshold with respect to the absolute differential TF activity in order to differentiate between TFs with low, medium and high signal, respectively (Fig. 2g and Suppl. Fig. 11).

**RNA-Seq processing for CLL data**

We used all ten available RNA-Seq samples from (Rendeiro et al., 2016) as well as available metadata (e.g., age, sex and condition). Initial quality control and adapter trimming (using *ILLUMINACLIP:Truseq-2.fa: 1:30:4:5:true TRAILING:3 MINLEN:20* instead) was performed as described in the ATAC-Seq pipeline. We then aligned the samples to hg19 using STAR (Dobin et al., 2013) with the parameters *--outFilterMultimapNmax 2 --quantMode GeneCounts* using the Gencode (Harrow et al., 2012) v19 annotation. We filtered 2 out of 10 samples due to data quality that we identified via PCA (data not shown, see also Suppl. Table 3). We finally employed *DESeq2* with the design formula “~ *condition*”, filtered genes with less than 5 reads on average in either one of the conditions, and identified those genes corresponding to the TFs from *HOCOMOCO*.

**Classification of TFs into activator and repressors for CLL data**

Firstly, we tried to classify TFs based on literature-mining, using *TRRUST*. However, we found that most TFs were classified almost equally often as activator and repressor, which makes it very difficult to determine an overall function for each TF (see Suppl. Fig. 12b). This is likely due to regulatory interactions being affected by multiple factors such as cell type, study conditions, highly studied TFs and different experimental conditions. Therefore, we decided to develop a cell-type specific data-driven approach to classify each TF into “activator”, “repressor” or “undetermined”, based on the overall effect on their putative TFBS. Our classifier is based on the assumption that increasing the level of an activating TF increases chromatin accessibility at its target sites while increasing the level of a repressing TF decreases it. For this, we calculated the Pearson correlation coefficients between the expression level of each TF and the ATAC-Seq signal of each putative TFBS across all individuals (see Suppl. Fig. 12a). If the median of the resulting correlation coefficients was positive we consider it an activator, if it is negative as repressor, and if it was not significantly different from the background we call it undetermined.

In detail, for each TFBS, we calculated a correlation median for (1) ATAC-Seq regions with at least one putative TFBS and (2) regions without a TFBS. The latter was used to estimate the noise level for the median correlations across all TFs. In particular, to distinguish real correlations from noise (i.e. activator/repressor from undetermined) we used the 5th (<0.040) and 95th (0.013) percentiles of the noise estimates as a threshold for activators and repressors respectively. The resulting classification of activators and repressors for positive and negative medians, respectively, was used to assess the level of coherence based on TF expression and their TF activity calculated by *diffTF.* We also summarised the fraction of the activators/repressors in certain TF cluster from RSAT clustering (Suppl. Fig. 14).

**Repressor/Activator classification validation I: Correlation of TF activity with target genes expression for CLL data**

For the CLL target analysis we used the same set of predicted TFBS as with *diffTF* and annotated each TFBS to the closest gene using the *ChIPseeker* (Yu et al., 2015) package in

R/Bioconductor (Huber et al., 2015) using a *hg19* annotation. TFBS located -2,000/+500 bp from the TSS were annotated as promoter TFBS, TFBS outside of -2,000/+500 but within +10kb/-10kb as enhancer, and the remaining few TFBS outside of +10kb/-10kb were discarded. Using expression data from RNA-seq as described above, we then calculated the mean and median log2 fold changes of the target genes of a given TF that were defined as having at least one TFBS of interest in the promoter regions. For the final representation in Fig. 5c, we used only TFs that had more than 200 and less than 1,500 unique target genes.

**Repressor/Activator classification validation II: Footprinting analysis for CLL data**

For the CLL footprinting analysis, we selected all 25 U-CLL samples and the 25 samples for M-CLL with the highest read counts, downsampled all to the sample with the lowest read count (~14.5 million) and merged all U-CLL and M-CLL samples. For all TFs that are expressed according to the RNA-Seq data, we then took the same set of TFBS as for *diffTF*, without TFBS that were overlapping between activators and repressors, and run the *dnase_to_javatreeview.py* script from the *pyDNase* library (Piper et al., 2015) for each condition to obtain base-specific Tn5 insertions matrices for each TFBS. These values were then normalized to the amount of reads in the consensus peakset, which was calculated using *featureCounts* from the *Subread* package with the parameters -p -B -d 0 -D 2000 -C -Q 10 -O -s 0. Finally, we removed peaks in blacklisted regions and potential artifacts (>1000 counts). For generating the genomic background for the openness, we binned the consensus peakset with a bin size of 200 bp and randomly selected 10,000 regions for which we then ran again the *dnase_to_javatreeview.py* script followed by subsequent normalization. The final matrices were clustered and visualised in Suppl. Fig. 13.

**Repressor/Activator classification validation III: *chromHMM* state enrichment**

For each TF deemed significant with *diffTF* that was also expressed according to the RNA-Seq data, we intersected all TFBS with the expanded 18-state model from the *chromHMM* data of the primary B-cells (Roadmap Epigenomics Consortium et al., 2015). For each TF, we then calculated the fraction of its TFBS overlapping each state. We then grouped the TFs into activators and repressors, visualized their respective distributions with boxplots, and statistically assessed the differences between activators and repressors using a Wilcoxon test.

**Comparing *diffTF* results with HOMER for CLL data**

We also assessed the detection power of *diffTF* compared to a standard motif enrichment analysis (Suppl. Fig. 18). For this, we generated a set of background sequences for each set of DA peaks in the two conditions, considering length and GC bias as features for the background generation. We then used *HOMER* (Heinz et al., 2010) to perform an enrichment analysis of motifs from *HOCOMOCO* on the DA peaks of each condition as foreground and respective two-fold *BiasAway* (Worsley Hunt et al., 2014) generated-sequences as background. We then correlated the *diffTF* results with the percentage of foreground sequences enriched for the given motif (HOMER output).

**Comparing *diffTF* with *chromVAR* results for CLL data**

We also compared our *diffTF* CLL results with *chromVAR (Schep et al., 2017)* (Suppl. Fig. 19). For this, we mimicked our *diffTF* analysis in the *chromVAR* framework as much as possible. First, we imported all TFBS that overlapped with our peaks and that were used in *diffTF* using the *getAnnotations* function in chromVAR. We then used the same consensus peakset as input as for *diffTF* (without resizing the peaks to maximize compatibility). We then derived fragment counts in paired-end mode for the same BAM files using the function *getCounts*, followed by GC correction using *addGCBias*, peak filtering using *filterPeaks* using the default arguments. To compute the expectations, we used the CLL group assignments into U-CLL and M-CLL while also normalizing counts, which yields within each group the average fraction of reads per peak in each sample. After calculating the deviations and deviation scores, we summarized the results per TF by calculating the difference of the means between U-CLL and M-CLL samples separately for both of them. Finally, we correlated the *chromVAR* results with the differential TF activity from *diffTF*.

Generally, as stated and quantified in the main text, *diffTF* and *chromVAR* correlate significantly (Pearson Suppl. Figure 19a-b) and agree for most TFs, but there are some at first sight discrepancies for TFs located in the second quadrant. They can likely be explained by the following methodological differences:

1. *chromVAR* compares only peaks and not individual TFBS, while *diffTF* computes a log2 fold-change for each TFBS. While peak log2 fold-change and corresponding TFBS log2 fold-change overall correlate highly (Pearson 0.91), TF-specific differences are in the range of -0.038 to 0.026 for expressed TFs in log2 fold-change units, which is

considerable (47.5 and 26%, respectively) given that the *diffTF* TF activity score is in the same scale and ranges from -0.08 to 0.10 (Suppl. Fig. 19c-d for two specific examples and E for a summary across TF). Overall, we observe that the mean log2 fold-change across all TFBS is 0.026, while for peaks, it is only 0.019.

2. Unlike *chromVAR, diffTF* compares each TF against the mean effect across all TF and therefore uses a relative rather than an absolute value. This explains most of the shift from TFs in Suppl. Fig. 19a in the left upper quadrant because overall, the mean log2 fold-change across all TFBS is slightly skewed to the positive. Thus, most *chromVAR* deviation values are consequently mostly positive (424 out of 640, ~66.3%), while *diffTF* TF activity values are mostly negative (420 out of 640, ~ 65.6%).
3. *chromVAR* never directly compares the two conditions with one another to compute log2 fold-changes, but only uses the conditions to compute a deviation that is based on the condition-specific expectation. Consequently, it provides a deviation value for each sample, which then have to be summarized accordingly. The recommended and used approach is to compute the mean deviation within each condition, the difference of which then mimics the differential TF activity as used in *diffTF*. However, the computation of the mean can be prone to outliers (Suppl. Fig. 19 F). *diffTF* does not directly suffer from this issue as it compares the two groups with one another to derive a log2 fold-change.

From all expressed TFs for which *chromVAR* and *diffTF* differ in their predicted direction of change, we checked in our literature review for the TFs that were deemed significant in *diffTF* which CLL condition they have been previously been associated with (Suppl. Table 2). Only 8 TFs are located in quadrant 2 (GCM1, NR4A3, NR1H2, NR1D1, PPARD, ESR1.A, NF2L1, ZBTB6). Out of these, only PPARD has been clearly associated with either of the two condition (M-CLL), as also predicted by diffTF. *chromVAR*, however, associates it more with U-CLL, although the deviation value is relatively small.

**HSC mouse data source (FACS sorting step)**

Single-cell suspensions of mouse bone marrow were erythrolysed, enriched for Kit expression (CD117 microbeads, Miltenyi Biotech) and stained with antibodies against surface markers: Lineage (B220-PECy5 (RA3-6B2, eBioscience), CD11b-PECy5 (M1/70, eBioscience), Ter119, PECy5 (TER-119, eBioscience), CD3e-PECy5 (145-2C11, eBioscience), Gr1-PECy5 (RB6-8C5, eBioscience)), Sca1-BV421 (D7, BD biosciences), cKit-AlexaFlour 780 (2B8, eBioscience),

CD150-APC (TC15-12F12.2, Biolegend), CD48-PE (HM48-1, eBiocience), CD16/32-PECy7 (93, eBioscience). The following combination of surface markers was used to define hematopoietic progenitor populations: Multipotent Progenitor (MPP) cells, Lin$^-$cKit$^+$Sca1$^+$CD150$^-$CD48$^+$; Granulocyte-Monocyte Progenitor (GMP) cells, Lin$^-$cKit$^+$Sca1$^-$CD16/32$^+$. Cells were sorted on *FACSAria III* (BD Biosciences) and analyzed using the *FlowJo* software (Tree Star inc.).

**HSC ATAC-Seq libraries generation**

ATAC-Seq libraries were generated as described previously (Buenrostro et al., 2013; Lara-Astiaso et al., 2014), with the following modifications. Briefly, 10.000 hematopoietic progenitor cells (MPPs or GMPs) freshly isolated from individual wild type mice were sorted into ice-cold FACS buffer (PBS + 2%FBS). The cells were pelleted using a swinging bucket centrifuge (500 x g, 10min, 4ºC) with settings for low acceleration/deceleration and washed once in ice-cold PBS. The cell pellets were resuspended in 50µl lysis buffer (10mM Tris-HCl pH 7.4, 10mM NaCl, 3mM $MgCl_2$, 0.1% Igepal CA-630) by gentle pipetting and immediately centrifuged one additional time (500 x g, 10min, 4ºC). The supernatant was discarded and the pellet containing released nuclei were resuspended gently in 25µl 1xTD buffer containing 1.25µl Tn5 transposase (Nextera sample preparation kit, Illumina). The transposition reaction was allowed to proceed for 45min at 37ºC whereafter DNA fragments were isolated using MinElute PCR purification columns (Qiagen) according to manufacturer's instruction.

To generate multiplex libraries, the transposed DNA were initially amplified for 5x PCR cycles using 2.5µl each of dual-index primers (Nextera index kit, Illumina) and 2.5µl PCR primer cocktail (PPC, Illumina) in a 25µl reaction volume of 1x KAPA HiFi hot-start ready-mix (Kapa BioSystems). The hot-start polymerase was activated prior to adding to the reaction mix by performing a brief pre-incubation step of 3min at 95ºC. The amplified fragments were size-selected with AMPure XP beads (0.5X) to remove fragments larger than 600bp and an aliquot was quantified to determine the optimal PCR cycle number to obtain 1/3 of maximum fluorescence intensity (Library quantification kit, Kapa Biosystems). Finally, PCR amplification was performed using the optimal number of cycles determined for each library (max. 18 cycles in total), size-selected with AMPure XP beads (0.5X) and eluted in resuspension buffer (Illumina). The size distribution of the libraries was evaluated on Bioanalyzer (Agilent) and sequenced on NextSeq 500 (Illumina) using 75bp paired-end sequencing with an average of 25 million reads per sample.

**GMP-MPP ATAC-Seq processing**

We run the ATAC-Seq pipeline described as above on the four wildtype GMP and four wildtype MPP samples with default parameters using the *mm10* genome. About 30% of the reads successfully passed all the filtering criteria of the pipeline, with the majority of the reads eliminated at the “remove duplicates” and “remove mitochondrial reads” steps. The fragment length distributions also showed the typical and expected pattern, with peaks at expected lengths (e.g, mono- and dinucleotides). We used GC correction as described above to remove technical GC sequencing biases.

**GMP-MPP *diffTF* analysis**

We run *diffTF* for the GMP-MPP dataset and compared the four GMP and MPP samples using default parameters unless otherwise specified. For identifying the consensus peak set we required a minimum of two samples that need to contain the peak, which resulted in 77,678 peaks. As parameters for the peaks and TF-specific analysis, we used only the stage of the HSC differentiation (GMP or MPP) in the design formula. Due to the small number of samples and therefore also possible permutations, we used the analytical approach as described in the Methods.

**GMP-MPP footprinting analysis**

For this footprinting analysis, we used all 8 ATAC-seq samples generated for MPP and GMP cell types and downsampled all to the sample with the lowest read counts (~8.3 million) and subsequently merged them by cell type. For all TFs from the RNA-Seq data that were significantly differentially expressed in DESeq2 (adj. p-value < 0.05), we then took the same set of TFBS as for *diffTF* and run the *dnase_to_javatreeview.py* script from the *pyDNase* library (Piper et al., 2015) for each condition to obtain base-specific Tn5 insertions matrices for each TFBS. These values were then normalized to the amount of reads in the consensus peakset, which was calculated using *featureCounts* from the *Subread* package with the parameters -p -B -d 0 -D 2000 -C -Q 10 -O -s 0. For generating the genomic background for the openness, we binned the consensus peakset with a bin size of 200 bp and randomly selected 10,000 regions for which we then ran again the *dnase_to_javatreeview.py* script followed by subsequent normalization. To generate the footprint plots for each cluster (Suppl. Fig. 17, left), we divided the value of Tn5 insertions at each bp to the mean value of Tn5 insertions in whole matrix for the specific cluster in order to enhance the differences between the clusters in the motif center and the surroundings.

**Software availability**

The *diffTF* pipeline as well as a detailed documentation are publicly available as a *Snakemake* workflow through *difftf.readthedocs.io*, from which the Github repository at *https://git.embl.de/grp-zaugg/diffTF* is also linked.

#### Supplementary Figures:

**Supplementary Figure 1.** Concept of the differential TF activity approach: Based on findings from chromatin QTL studies that have reported changes in chromatin often coincide with SNPs disrupting TF binding sites (TFBS), we propose to detect global differences in TF activity by investigating chromatin accessibility (or modifications) in close proximity to their TFBS.

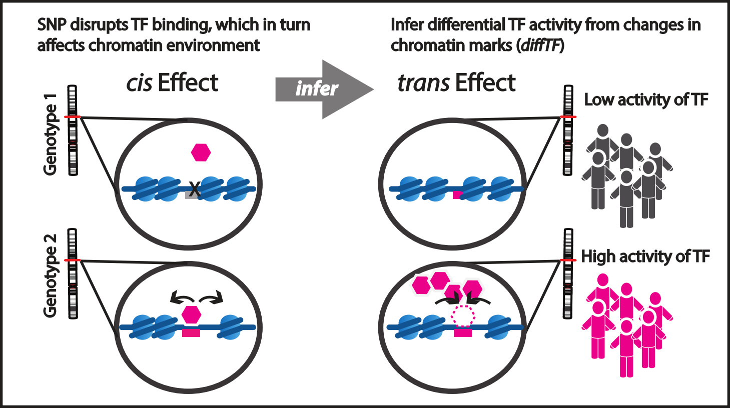
SNP disrupts TF binding, which in turn affects chromatin environment
Infer differential TF activity from changes in chromatin marks (diffTF)
cis Effect
infer
trans Effect
Genotype 1
Genotype 2
X
Low activity of TF
High activity of TF

**Supplementary Figure 2.** Summary workflow of the ATAC-Seq (top) and *diffTF* (bottom) pipelines.

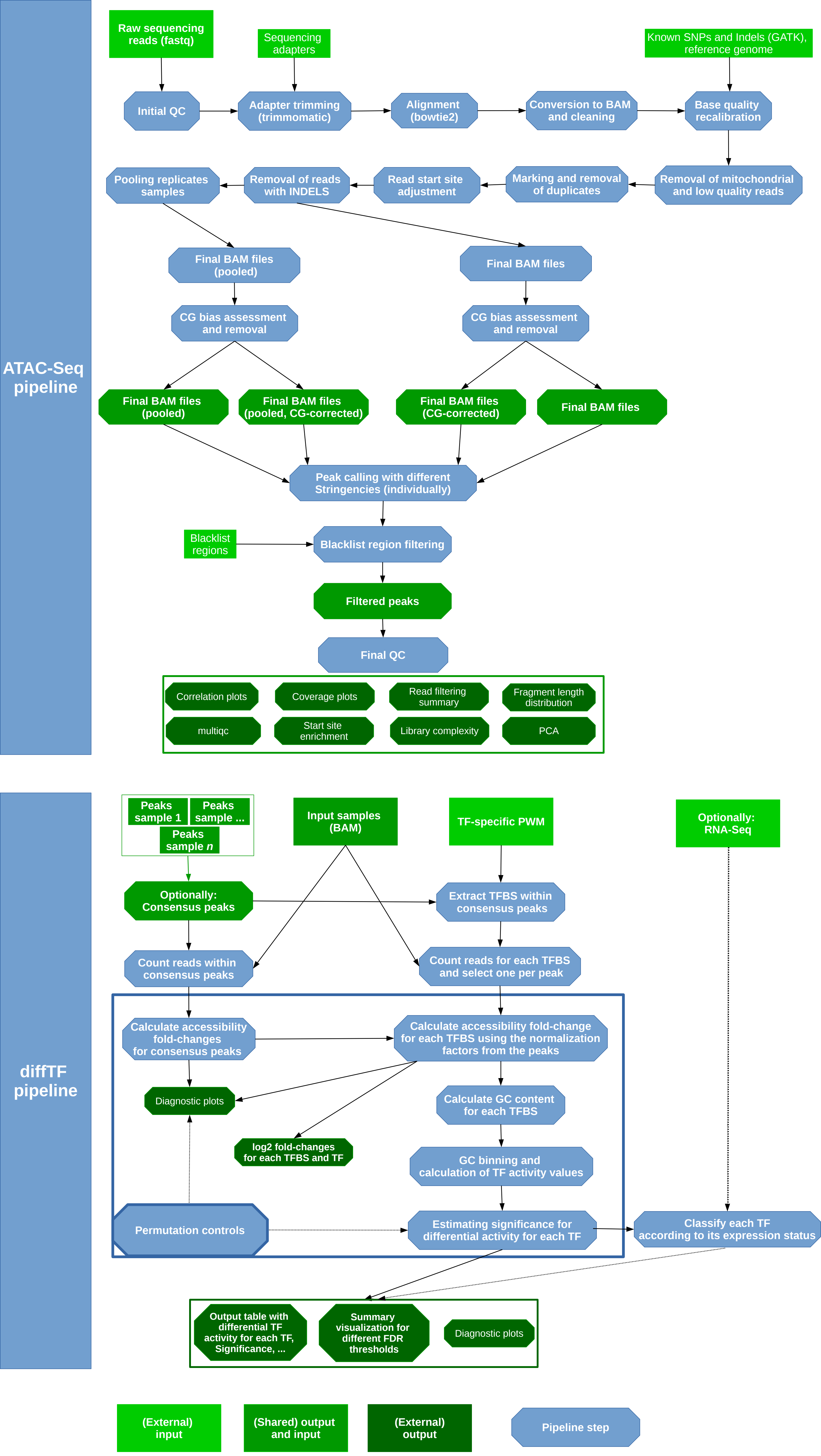

ATAC-Seq pipeline
Raw sequencing reads (fastq)
Sequencing adapters
Known SNPs and Indels (GATK), reference genome
Initial QC
Adapter trimming (trimmomatic)
Alignment (bowtie2)
Conversion to BAM and cleaning
Base quality recalibration
Removal of mitochondrial and low quality reads
Marking and removal of duplicates
Read start site adjustment
Removal of reads with INDELS
Pooling replicates samples
Final BAM files (pooled)
Final BAM files
CG bias assessment and removal
CG bias assessment and removal
Final BAM files (pooled)
Final BAM files (pooled, CG-corrected)
Final BAM files (CG-corrected)
Final BAM files
Peak calling with different Stringencies (individually)
Blacklist regions
Blacklist region filtering
Filtered peaks
Final QC
Correlation plots
Coverage plots
Read filtering summary
Fragment length distribution
multiqc
Start site enrichment
Library complexity
PCA
diffTF pipeline
Peaks sample 1
Peaks sample ...
Peaks sample n
Input samples (BAM)
TF-specific PWM
Optionally: RNA-Seq
Optionally: Consensus peaks
Extract TFBS within consensus peaks
Count reads within consensus peaks
Count reads for each TFBS and select one per peak
Calculate accessibility fold-changes for consensus peaks
Calculate accessibility fold-change for each TFBS using the normalization factors from the peaks
Diagnostic plots
Calculate GC content for each TFBS
log2 fold-changes for each TFBS and TF
GC binning and calculation of TF activity values
Permutation controls
Estimating significance for differential activity for each TF
Classify each TF according to its expression status
Output table with differential TF activity for each TF, Significance, ...
Summary visualization for different FDR thresholds
Diagnostic plots
(External) input
(Shared) output and input
(External) output
Pipeline step

**Supplementary Figure 3.** Diagnostic plot for how the differential TF activity is calculated in *diffTF*. As example, IRF8 (A) and ZFHX3 (B) from the CLL dataset are shown. The top rows show the mean difference values across the different CG bins, with points being sized according to their weight (that is, the number of binding sites). In the example for IRF, the biggest contribution comes from the 40 and 50 % bins. The final measure, the differential TF activity, is also indicated as a dashed line. The lower panel shows the log2 fold-change distributions of the bin-specific foreground (the specific TF only) and background (all other TFs), respectively. Empty bins indicate that not enough binding sites (<5) were available.

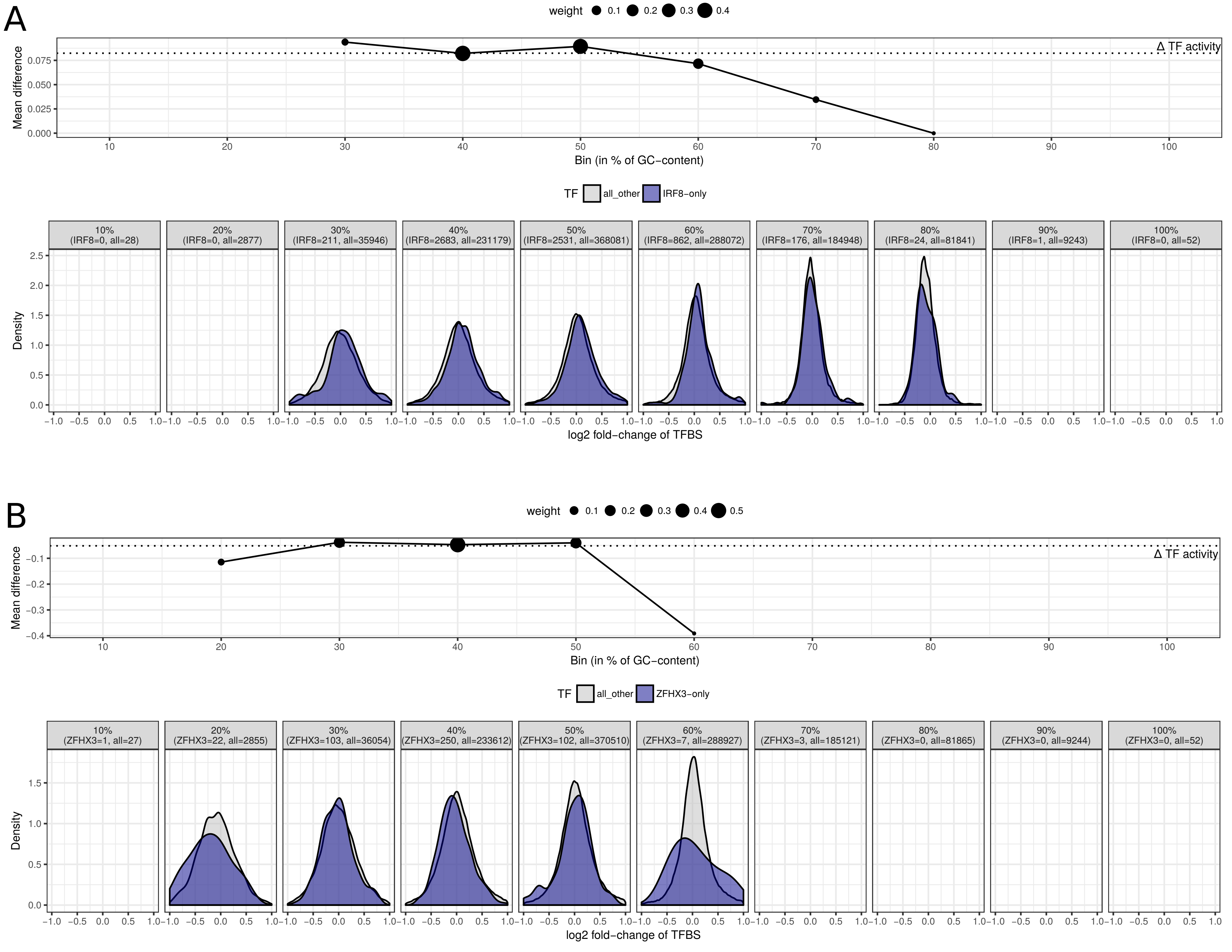

A
weight 0.1 0.2 0.3 0.4
Δ TF activity
Mean difference
Bin (in % of GC−content)
TF all_other IRF8−only
10% (IRF8=0, all=28)
20% (IRF8=0, all=2877)
30% (IRF8=211, all=35946)
40% (IRF8=2683, all=231179)
50% (IRF8=2531, all=368081)
60% (IRF8=862, all=288072)
70% (IRF8=176, all=184948)
80% (IRF8=24, all=81841)
90% (IRF8=1, all=9243)
100% (IRF8=0, all=52)
Density
log2 fold−change of TFBS
B
weight 0.1 0.2 0.3 0.4 0.5
Δ TF activity
Mean difference
Bin (in % of GC−content)
TF all_other ZFHX3−only
10% (ZFHX3=1, all=27)
20% (ZFHX3=22, all=2855)
30% (ZFHX3=103, all=36054)
40% (ZFHX3=250, all=233612)
50% (ZFHX3=102, all=370510)
60% (ZFHX3=7, all=288927)
70% (ZFHX3=3, all=185121)
80% (ZFHX3=0, all=81865)
90% (ZFHX3=0, all=9244)
100% (ZFHX3=0, all=52)
Density
log2 fold−change of TFBS

**Supplementary Figure 4.** Diagnostic plots for the empirical, permutation-based (A-E) for the CLL dataset as well as the analytical approach using the GMP/MPP dataset (F-I) in *diffTF*. (A) - (C) TF-specific permutation density plots for two significant TFs (B: EGR3, D: IRF2) and one non-significant one (C: E4F1). The real value is indicated as a red line. (D) Black line: Density distribution of the TF activity across all 640 TF and (i) all 1000 permutations and (N = 640000 Bandwidth = 0.0006962, black) and (ii) the real, non-permuted data (red). Expectedly, the permuted density centers around 0, whereas the density for the real values shows a clear shift towards negative values. (E) and (H) Scatterplot for the number of TFBS and the raw p-value (-log10 transformed) for the permutation-based (E) and the analytical approach (H). See text for details. (F) - (G) raw p-value (-log10 transformed) and T statistic (F and G, respectively) for each GC bin and across all bins ("all") for 3 selected TFs. The size of the dots indicates the number of TFBS in the bin and therefore its weight. (I). QQ plot of expected vs. observed p-values from the analytical approach (-log10 transformed). Red line: p-values for the real data. Black line: p-values for shuffled data (see Supplementary methods).

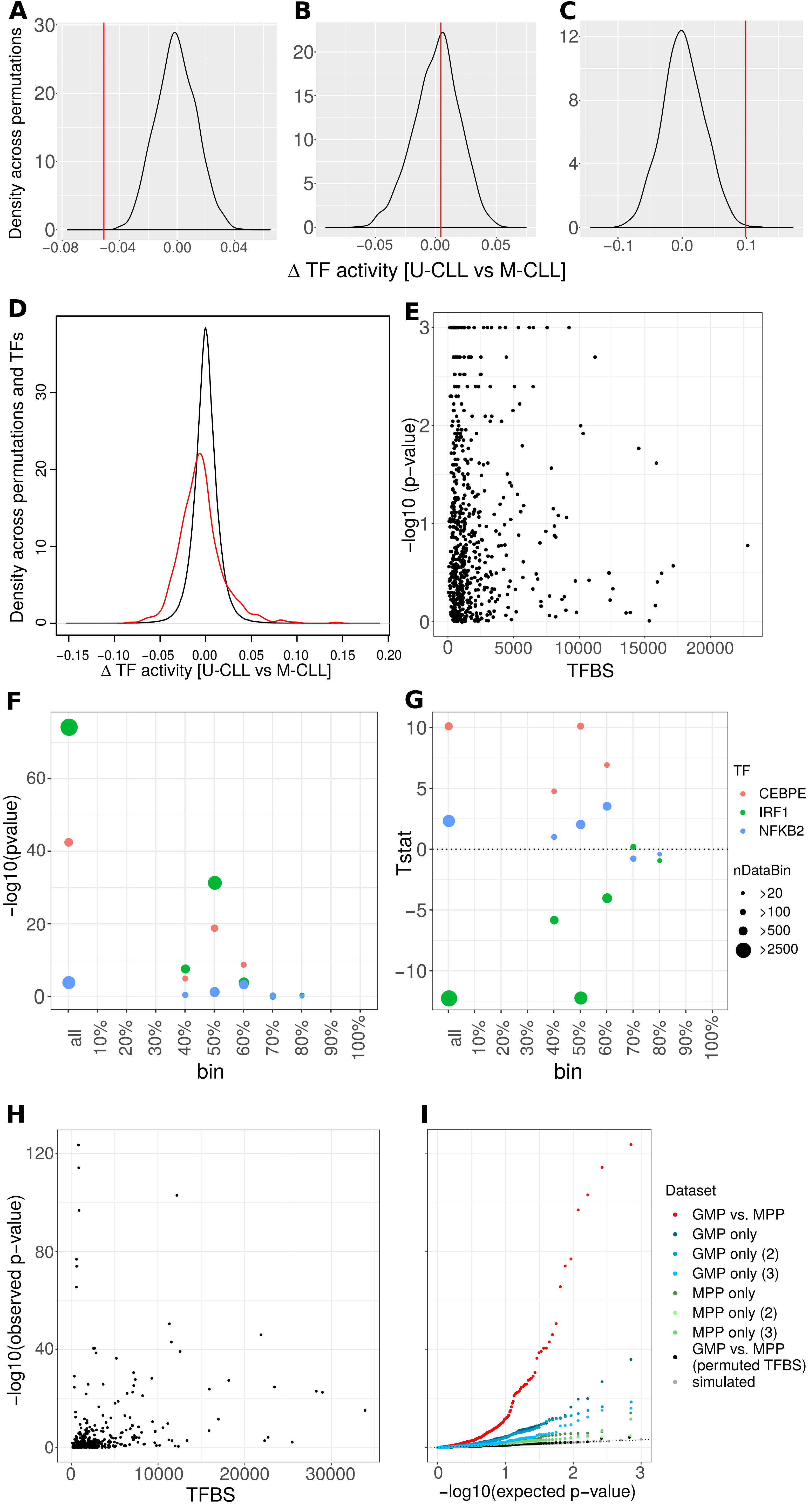
A
B
C
Density across permutations
Δ TF activity [U-CLL vs M-CLL]
D
Density across permutations and TFs
Δ TF activity [U-CLL vs M-CLL]
E
−log10 (p−value)
TFBS
F
-log10(pvalue)
bin
G
Tstat
bin
TF
CEBPE
IRF1
NFKB2
nDataBin
>20
>100
>500
>2500
H
−log10(observed p−value)
TFBS
I
−log10(expected p−value)
Dataset
GMP vs. MPP
GMP only
GMP only (2)
GMP only (3)
MPP only
MPP only (2)
MPP only (3)
GMP vs. MPP (permuted TFBS)
simulated

**Supplementary Figure 5.** Diagnostic plots for the *limma* analysis for the consensus peakset and selected TF-specific log2 fold-change distribution comparisons along with the consensus peakset for the CLL data. (A) and (B): MA plots for $\alpha$ = 0.1 for the mean of normalized counts against the log2 fold-change. (B) Empirical cumulative distribution function of the mean log counts for all samples after normalization. (C) and (D) Log2 fold-change density distribution and boxplots for three exemplary TFs that are either differentially active in mutated (CENPB), unmutated (IRF2) or in none of both (ETV7), as exemplified by their median log2 fold-changes.

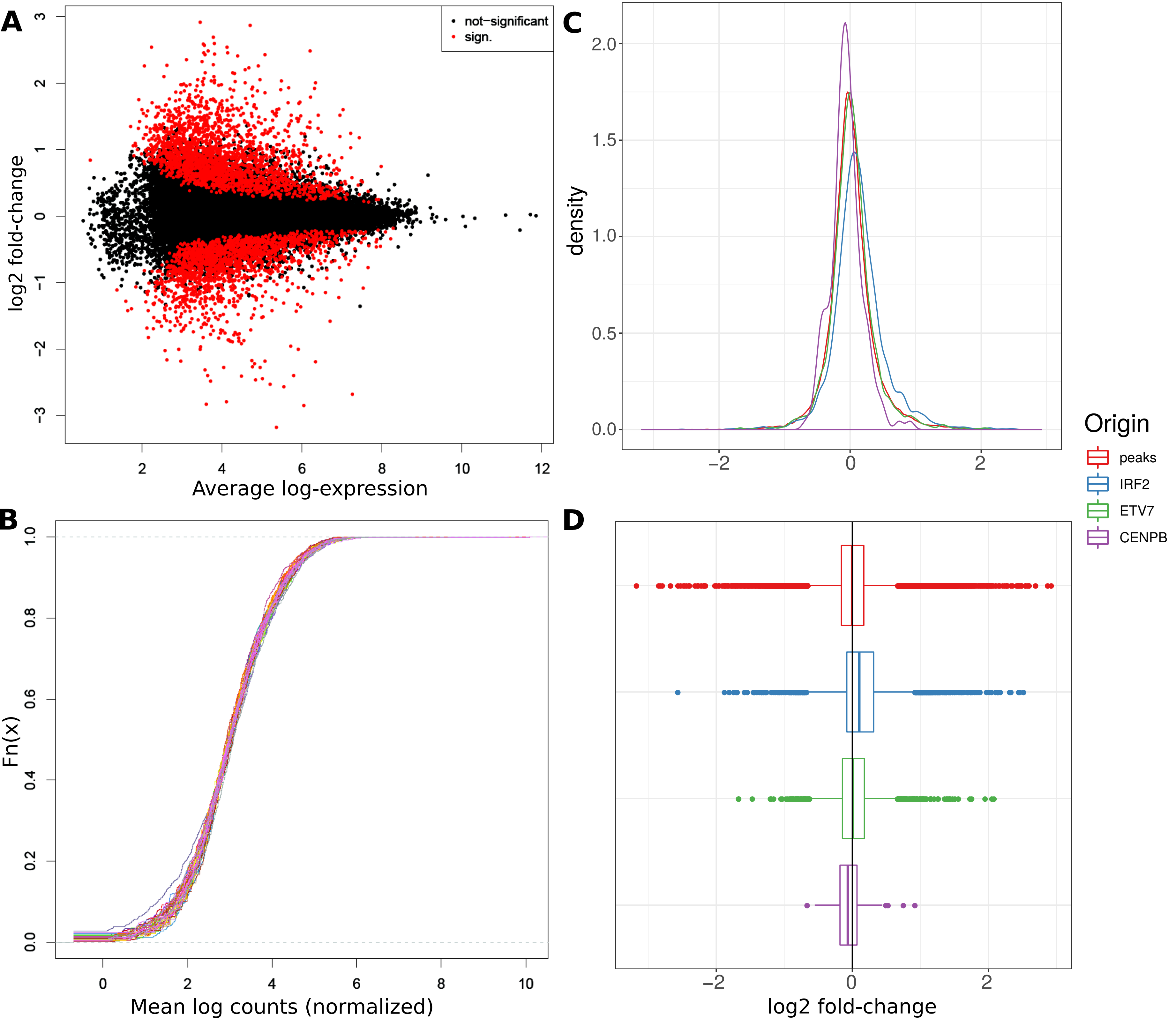
A
B
C
D
not−significant
sign.
log2 fold-change
Average log-expression
Fn(x)
Mean log counts (normalized)
density
Origin
peaks
IRF2
ETV7
CENPB
log2 fold-change

**Supplementary Figure 6.** Various statistics from our ATAC-Seq pipeline for the CLL data. (A) and (B): Summary statistics of the distribution of the percentage (A) and total number of reads (B) that passed the various filtering steps, respectively. Note that the various filtering criteria have been applied before any GC correction has been performed. (C) to (E): Individual statistics for one randomly chosen sample. (C): Fragment length distribution. (D) and (E): Estimated GC bias for 300 bp regions and the corresponding log2 ratios of observed/expected read counts (normalized) before (D) and after (E) applying the GC correction. The GC correction has been assessed with computeGCBias from *deepTools*. We used the options *--effectiveGenomeSize 2750000000* and *--blackListFileName* to exclude blacklisted regions.

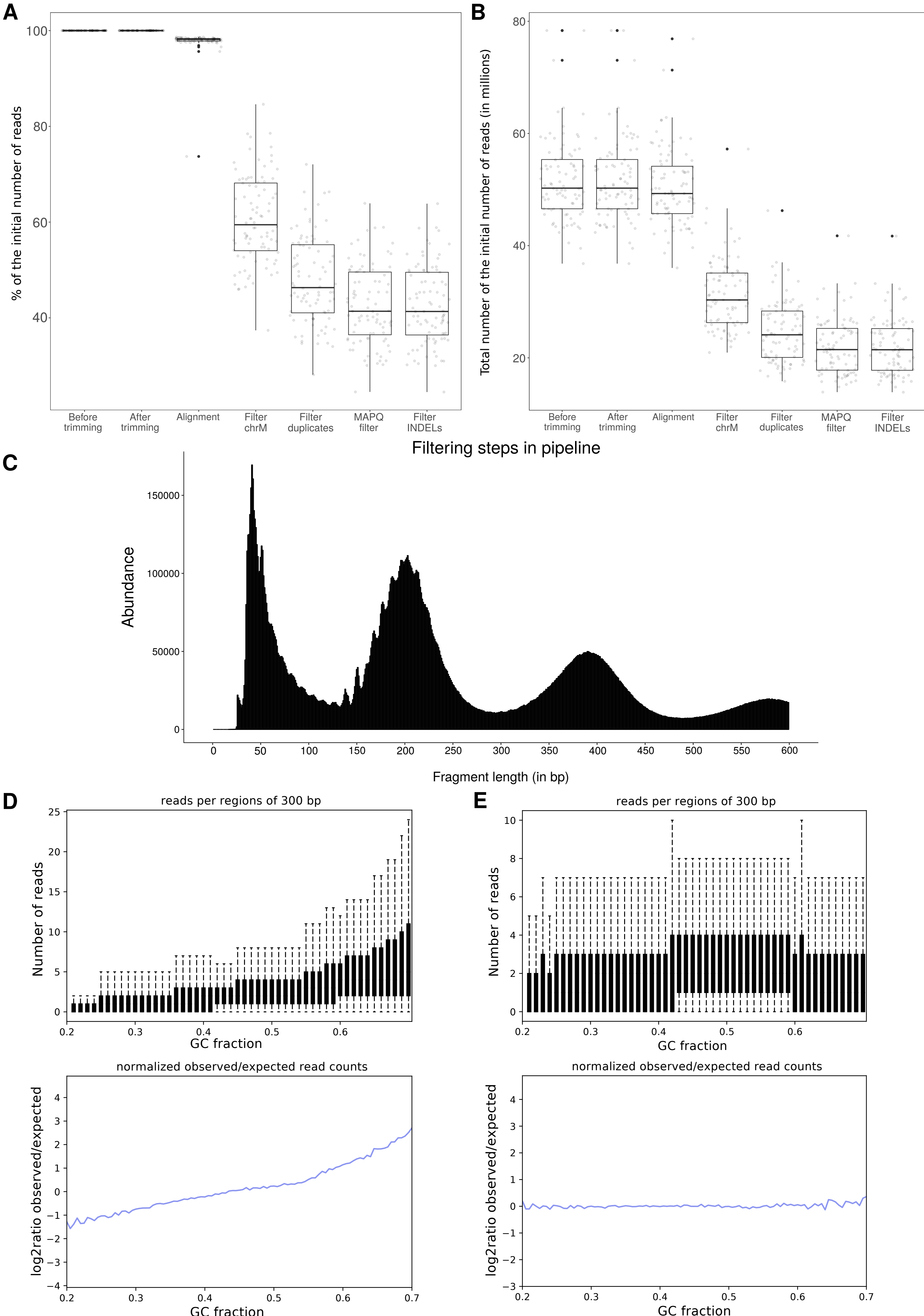

A
% of the initial number of reads
B
Total number of the initial number of reads (in millions)
Before trimming
After trimming
Alignment
Filter chrM
Filter duplicates
MAPQ filter
Filter INDELs
Filtering steps in pipeline
C
Abundance
Fragment length (in bp)
D
reads per regions of 300 bp
Number of reads
GC fraction
normalized observed/expected read counts
log2ratio observed/expected
GC fraction
E
reads per regions of 300 bp
Number of reads
GC fraction
normalized observed/expected read counts
log2ratio observed/expected
GC fraction

**Supplementary Figure 7.** PCA plots for all available metadata for the CLL data based on the consensus peaks. See Methods for details. (A) Treatment. (B) Condition. (C) IGVH homology. (D) Gender. (E) Patient age at data collection. (F) Patient age at diagnosis. Raw read counts have been produced with *multiBamSummary* from *deepTools* in BED-file mode using the option *--outRawCounts* for all consensus peaks. We then used *DESeq2* with the design "~ Treatment + Condition", variance-stabilized the data via the function *varianceStabilizingTransformation* using *BLIND=TRUE* and plotted the PCA data for all peaks with *plotPCA* for all available metadata.

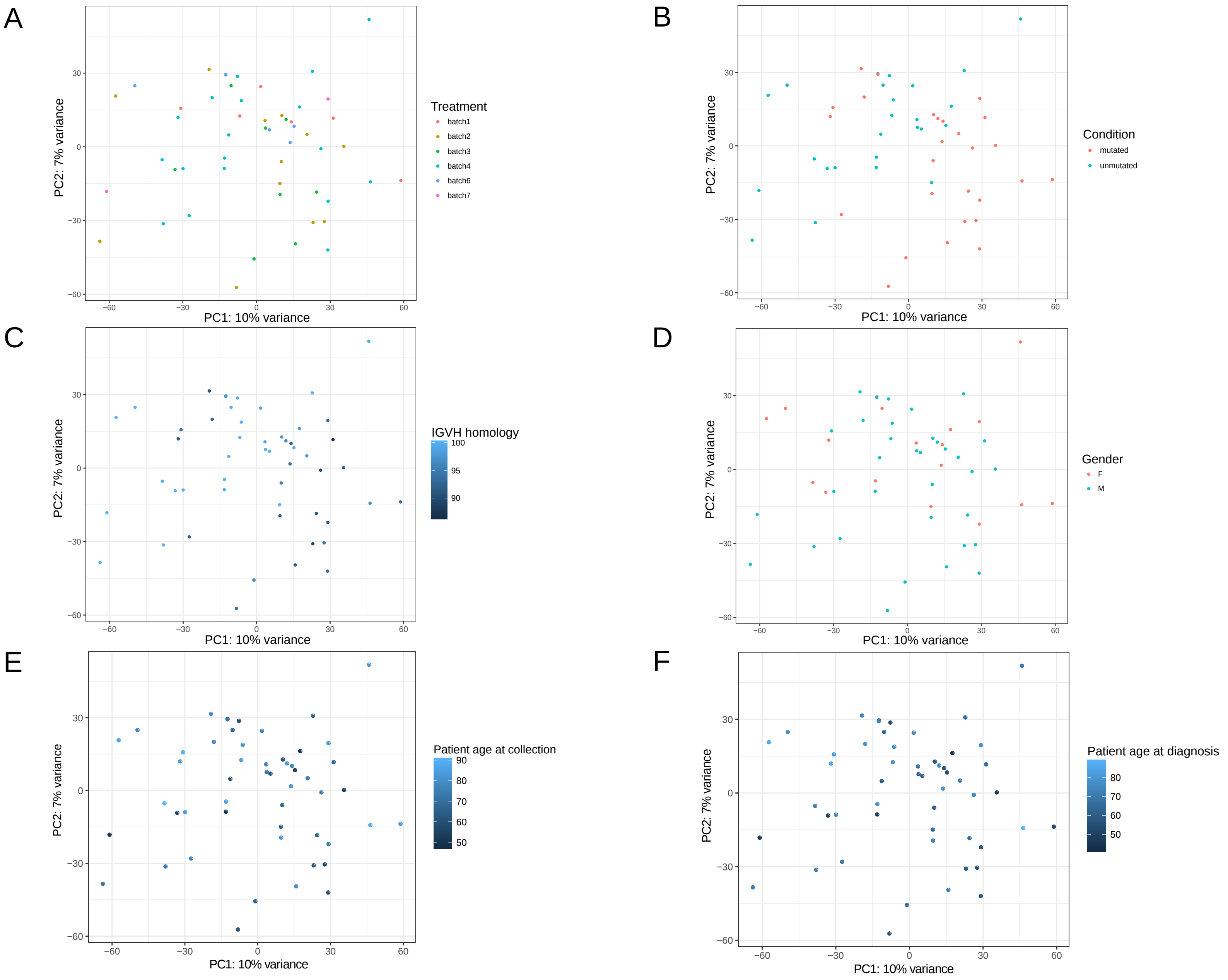
A
B
C
D
E
F
PC1: 10% variance
PC2: 7% variance
Treatment
batch1
batch2
batch3
batch4
batch6
batch7
Condition
mutated
unmutated
IGVH homology
Gender
F
M
Patient age at collection
Patient age at diagnosis

**Supplementary Figure 8.** Additional scatterplots of the differential TF activity related to *ReMap*. (A) All TFBS against only TFBS not intersecting with *ReMap* data. (B) Only TFBS not intersecting with *ReMap* against only TFBS intersecting with *ReMap*.

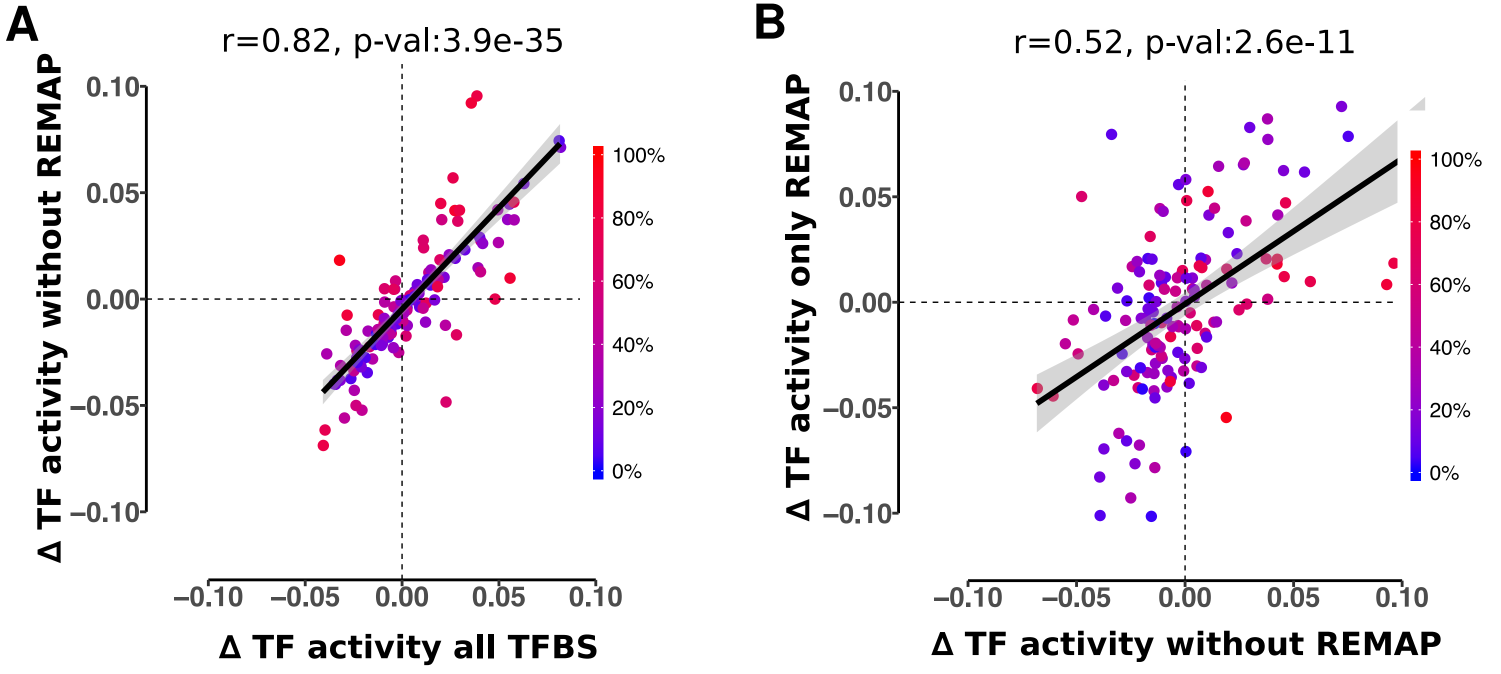
A
r=0.82, p-val:3.9e-35
Δ TF activity without REMAP
Δ TF activity all TFBS
0.10
0.05
0.00
−0.05
−0.10
−0.10
−0.05
0.00
0.05
0.10
100%
80%
60%
40%
20%
0%
B
r=0.52, p-val:2.6e-11
Δ TF activity only REMAP
Δ TF activity without REMAP
0.10
0.05
0.00
−0.05
−0.10
−0.10
−0.05
0.00
0.05
0.10
100%
80%
60%
40%
20%
0%

**Supplementary Figure 9.** Comparison of the effect of the nucleotide composition on the predicted binding sites for *PWMScan* and subsequent results for *diffTF*. (A) Scatterplot of the differential TF activity from all TFs for two different *diffTF* analyses are shown. Each point represents one TF. The *diffTF* analysis on the x-axis is based on the TFBS predicted by *PWMScan* with the nucleotide composition estimated from the full genome (0.29;0.21;0.21;0.29 for A, C, G, and T, respectively), while the y-axis results are based on the TFBS with the nucleotide composition estimated from the peaks only (0.27;0.23;0.23;0.27). Colors represent significance (FDR < 0.1) in analogy to Figure 2: white - not significant in either analysis; light green and light blue - significant only for the analysis from the x-axis or y-axis, respectively; purple - significant for both analyses. Spearman correlation was computed across all 640 TFs. (B) TFBS similarity between predicted TFBS using the genome-wide background versus the peak-specific one. Each point summarizes all TFBS from one TF. The x-axis shows the overall GC content of all TFBS from a particular TF overlapping with the peaks based on the genome-wide nucleotide composition prediction. The y-axis shows the similarity of all TFBS with the predicted TFBS based on the peak-specific nucleotide composition prediction as measured by the Jaccard index using *bedtools jaccard*.

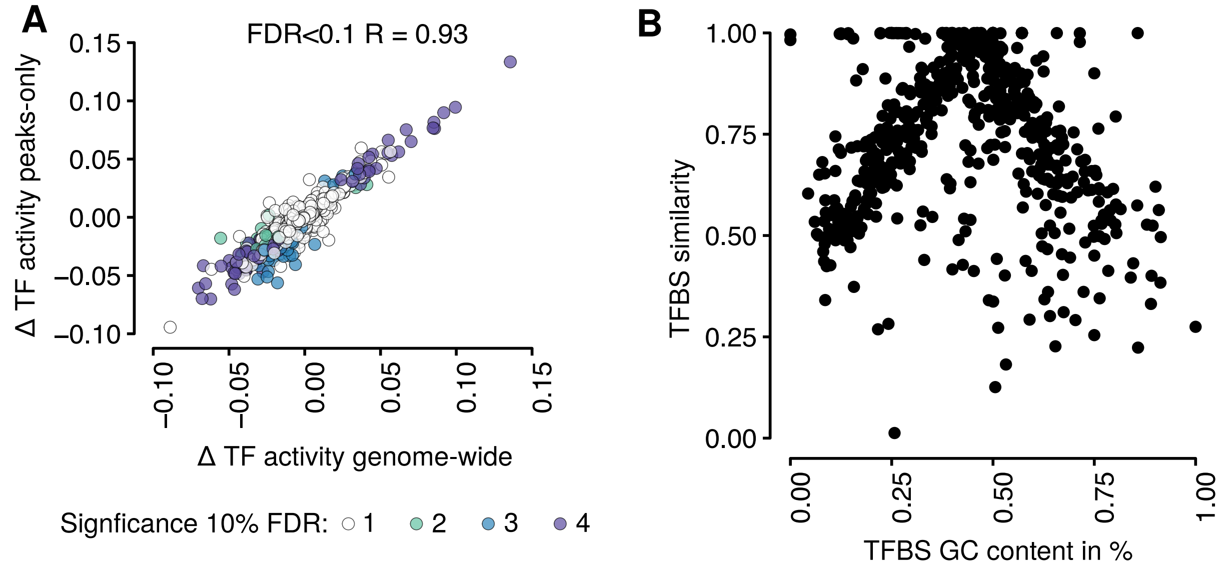
A
FDR<0.1 R = 0.93
Δ TF activity peaks-only
Δ TF activity genome-wide
0.15
0.10
0.05
0.00
−0.05
−0.10
Signficance 10% FDR: 1 2 3 4
B
TFBS similarity
TFBS GC content in %
1.00
0.75
0.50
0.25
0.00

**Supplementary Figure 10.** *diffTF* results for different motif extension sizes (0 to 600) for the CLL dataset. (A) Boxplot of the absolute differential TF activity across all TFs classified as either activator or repressor. (B) Heatmap of the differential TF activity across all TF, with additional metadata from the original analysis that used an extension size of 100.

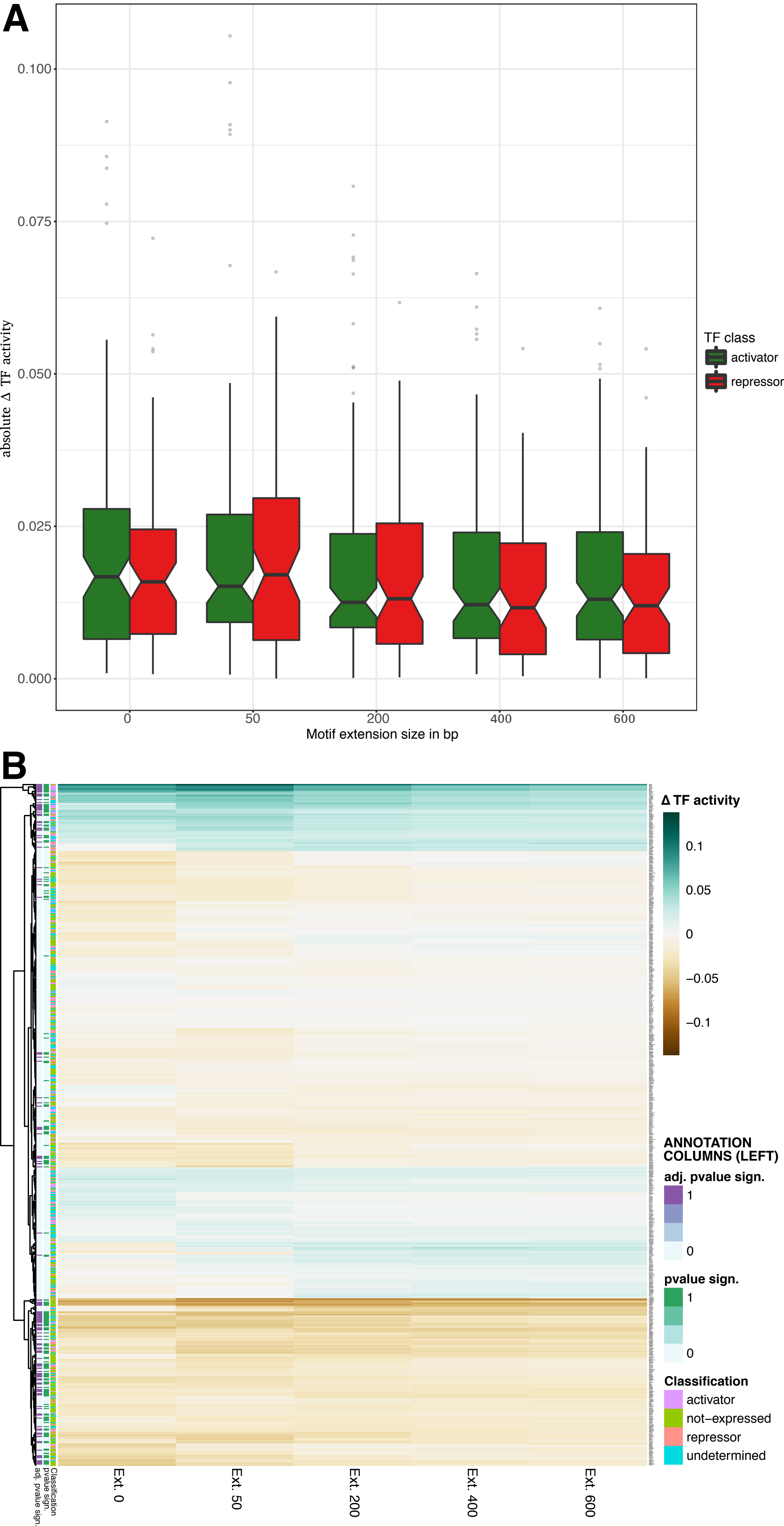
A
0.100
0.075
0.050
0.025
0.000
absolute Δ TF activity
0
50
200
400
600
Motif extension size in bp
TF class
activator
repressor
B
Δ TF activity
0.1
0.05
0
−0.05
−0.1
ANNOTATION COLUMNS (LEFT)
adj. pvalue sign.
1
0
pvalue sign.
1
0
Classification
activator
not−expressed
repressor
undetermined
Ext. 0
Ext. 50
Ext. 200
Ext. 400
Ext. 600
Classification
pvalue sign.
adj. pvalue sign.

**Supplementary Figure 11.** Further subsampling results. Same as Figure 2g, except that the fractions are shown only for those TFs that were not deemed significant with the full data.

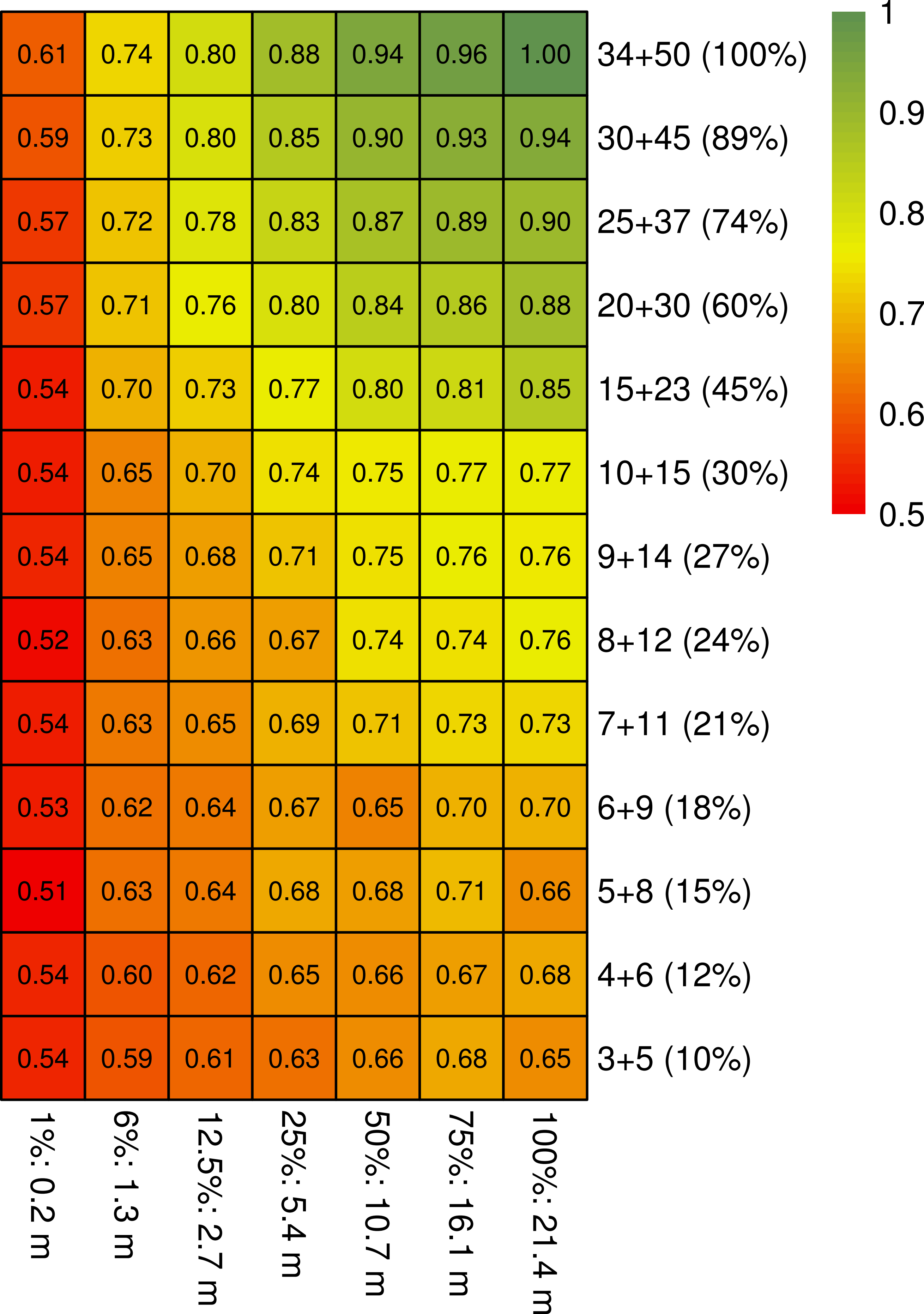

0.61 0.74 0.80 0.88 0.94 0.96 1.00
34+50 (100%)
0.59 0.73 0.80 0.85 0.90 0.93 0.94
30+45 (89%)
0.57 0.72 0.78 0.83 0.87 0.89 0.90
25+37 (74%)
0.57 0.71 0.76 0.80 0.84 0.86 0.88
20+30 (60%)
0.54 0.70 0.73 0.77 0.80 0.81 0.85
15+23 (45%)
0.54 0.65 0.70 0.74 0.75 0.77 0.77
10+15 (30%)
0.54 0.65 0.68 0.71 0.75 0.76 0.76
9+14 (27%)
0.52 0.63 0.66 0.67 0.74 0.74 0.76
8+12 (24%)
0.54 0.63 0.65 0.69 0.71 0.73 0.73
7+11 (21%)
0.53 0.62 0.64 0.67 0.65 0.70 0.70
6+9 (18%)
0.51 0.63 0.64 0.68 0.68 0.71 0.66
5+8 (15%)
0.54 0.60 0.62 0.65 0.66 0.67 0.68
4+6 (12%)
0.54 0.59 0.61 0.63 0.66 0.68 0.65
3+5 (10%)
1%: 0.2 m
6%: 1.3 m
12.5%: 2.7 m
25%: 5.4 m
50%: 10.7 m
75%: 16.1 m
100%: 21.4 m
1
0.9
0.8
0.7
0.6
0.5

**Supplementary Figure 12.** Classification of TFs into activators and repressors. (A) TF median correlations order from positive (top) to negative (bottom). TFs were classified as activator (green) or repressor (red) when the correlation with their putative binding sites was outside the 95th (green vertical line) and 5th (red vertical line) percentile from the distribution of all correlations at non-putative TFBS, respectively. (B) Barplot of the number of studies detected by text-mining for expressed TFs in CLL whose transcriptional activity on studied target gene was classified as activator (green bars) or repressor (red).

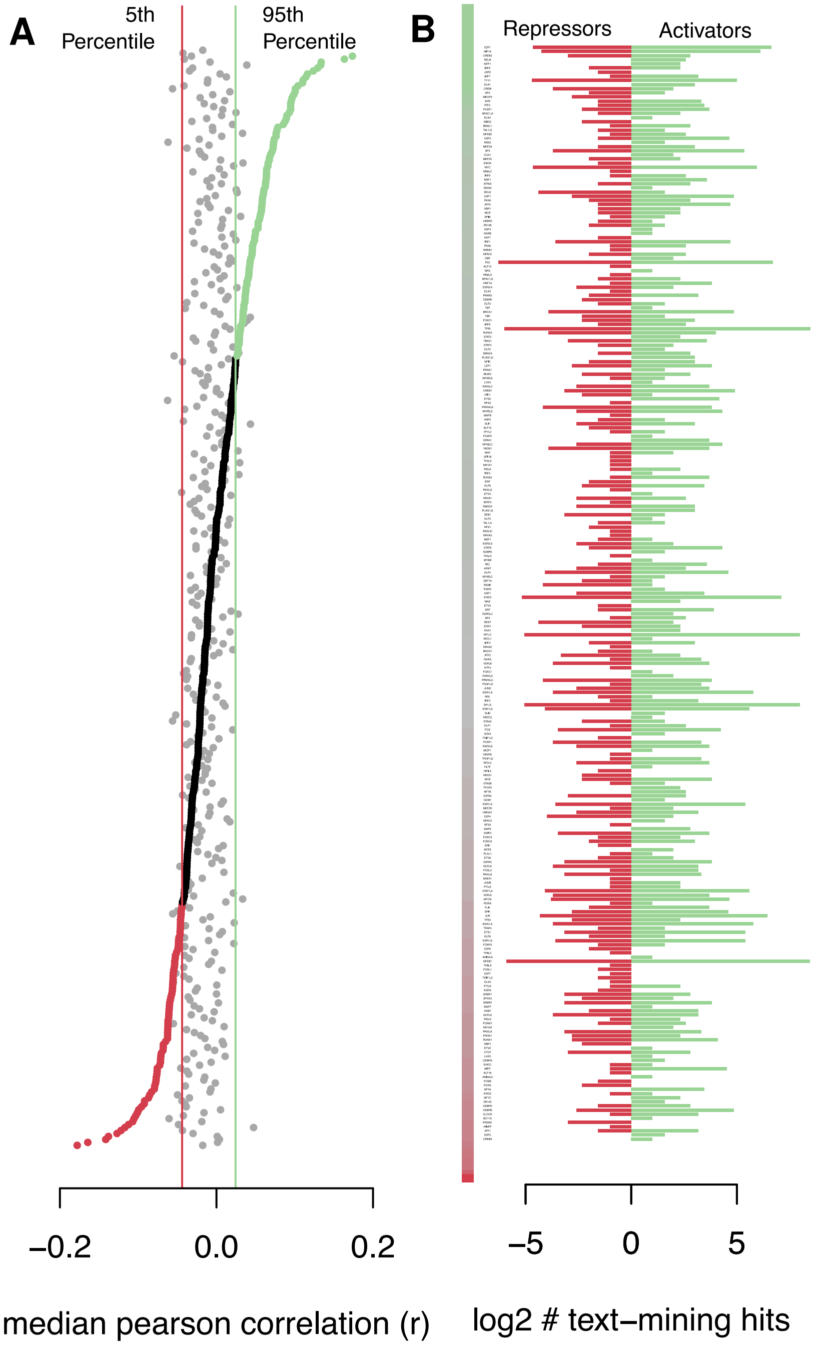
A
5th Percentile
95th Percentile
−0.2
0.0
0.2
median pearson correlation (r)
B
Repressors
Activators
−5
0
5
log2 # text−mining hits

**Supplementary Figure 13.** Cluster heatmap for the footprints for CLL. The heatmap shows TF footprint values for all significant TFs from *diffTF* (adj. p-value < 0.05), based on the normalised Tn5 insertions for each bp (+/-100bp from motif center). Colors represent footprint strength, while white denotes the value of the genomic background in the consensus peakset. Clusters were defined using hierarchical clustering with the *ward.D2* method.

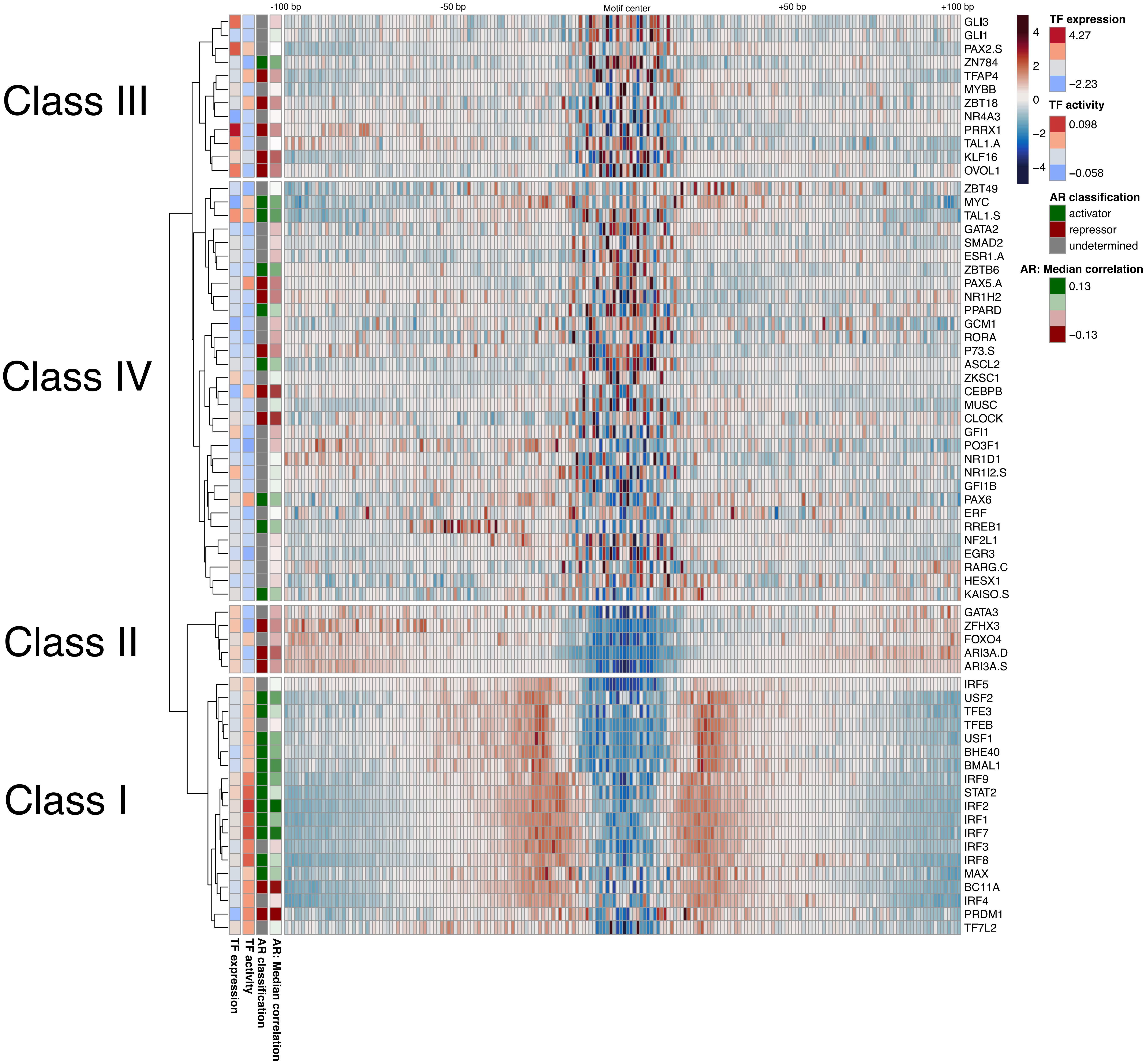

-100 bp
-50 bp
Motif center
+50 bp
+100 bp
Class III
Class IV
Class II
Class I
GLI3
GLI1
PAX2.S
ZN784
TFAP4
MYBB
ZBT18
NR4A3
PRRX1
TAL1.A
KLF16
OVOL1
ZBT49
MYC
TAL1.S
GATA2
SMAD2
ESR1.A
ZBTB6
PAX5.A
NR1H2
PPARD
GCM1
RORA
P73.S
ASCL2
ZKSC1
CEBPB
MUSC
CLOCK
GFI1
PO3F1
NR1D1
NR1I2.S
GFI1B
PAX6
ERF
RREB1
NF2L1
EGR3
RARG.C
HESX1
KAISO.S
GATA3
ZFHX3
FOXO4
ARI3A.D
ARI3A.S
IRF5
USF2
TFE3
TFEB
USF1
BHE40
BMAL1
IRF9
STAT2
IRF2
IRF1
IRF7
IRF3
IRF8
MAX
BC11A
IRF4
PRDM1
TF7L2
TF expression
TF activity
AR classification
AR: Median correlation
4
2
0
−2
−4
TF expression
4.27
−2.23
TF activity
0.098
−0.058
AR classification
activator
repressor
undetermined
AR: Median correlation
0.13
−0.13

**Supplementary Figure 14.** Functional clustering of TFs based on the similarity of their PWMs. The clustering is identical to Fig. 3, except that at the right side, the distribution of activators, repressors, and undetermined TFs is displayed for each cluster. TFs are also colored by their classification. See Fig. 3 for more details.

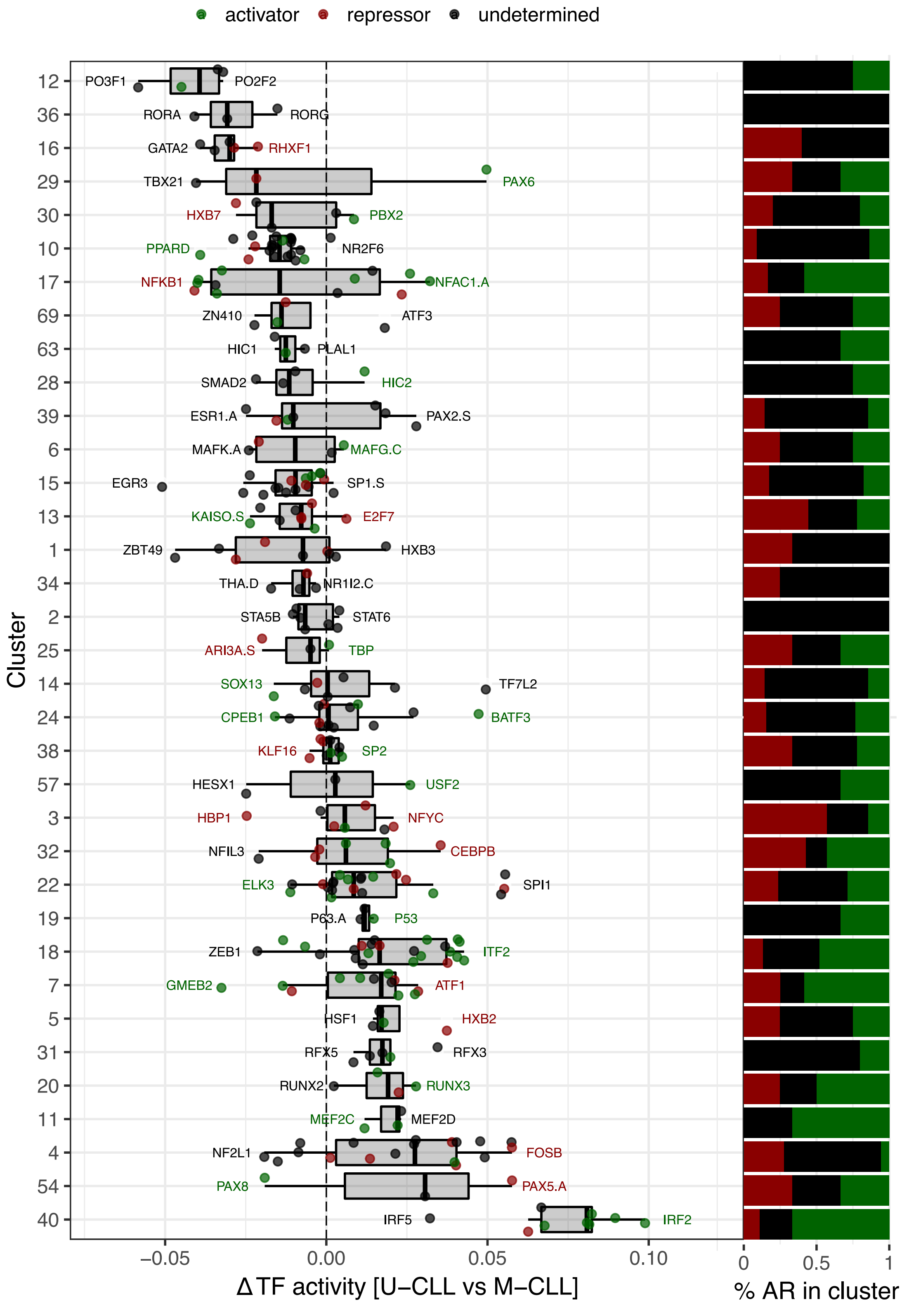
activator
repressor
undetermined
Cluster
12
36
16
29
30
10
17
69
63
28
39
6
15
13
1
34
2
25
14
24
38
57
3
32
22
19
18
7
5
31
20
11
4
54
40
PO3F1
PO2F2
RORA
RORG
GATA2
RHXF1
TBX21
PAX6
HXB7
PBX2
PPARD
NR2F6
NFKB1
NFAC1.A
ZN410
ATF3
HIC1
PLAL1
SMAD2
HIC2
ESR1.A
PAX2.S
MAFK.A
MAFG.C
EGR3
SP1.S
KAISO.S
E2F7
ZBT49
HXB3
THA.D
NR1I2.C
STA5B
STAT6
ARI3A.S
TBP
SOX13
TF7L2
CPEB1
BATF3
KLF16
SP2
HESX1
USF2
HBP1
NFYC
NFIL3
CEBPB
ELK3
SPI1
P63.A
P53
ZEB1
ITF2
GMEB2
ATF1
HSF1
HXB2
RFX5
RFX3
RUNX2
RUNX3
MEF2C
MEF2D
NF2L1
FOSB
PAX8
PAX5.A
IRF5
IRF2
−0.05
0.00
0.05
0.10
ΔTF activity [U−CLL vs M−CLL]
0
0.5
1
% AR in cluster

**Supplementary Figure 15.** Overlap of TFBS for activators and repressors with the 18-state *chromHMM* model for primary B-cells. For each state, we plot the distribution of the fraction of TFBS overlapping with the particular state as compared to the overall number of TFBS per TF. Boxplots are done separately for activators (green) and repressors (red), see text for details. In addition, we quantified the differences between activators and repressors using a Wilcoxon test (see the raw p-value above the boxplots).

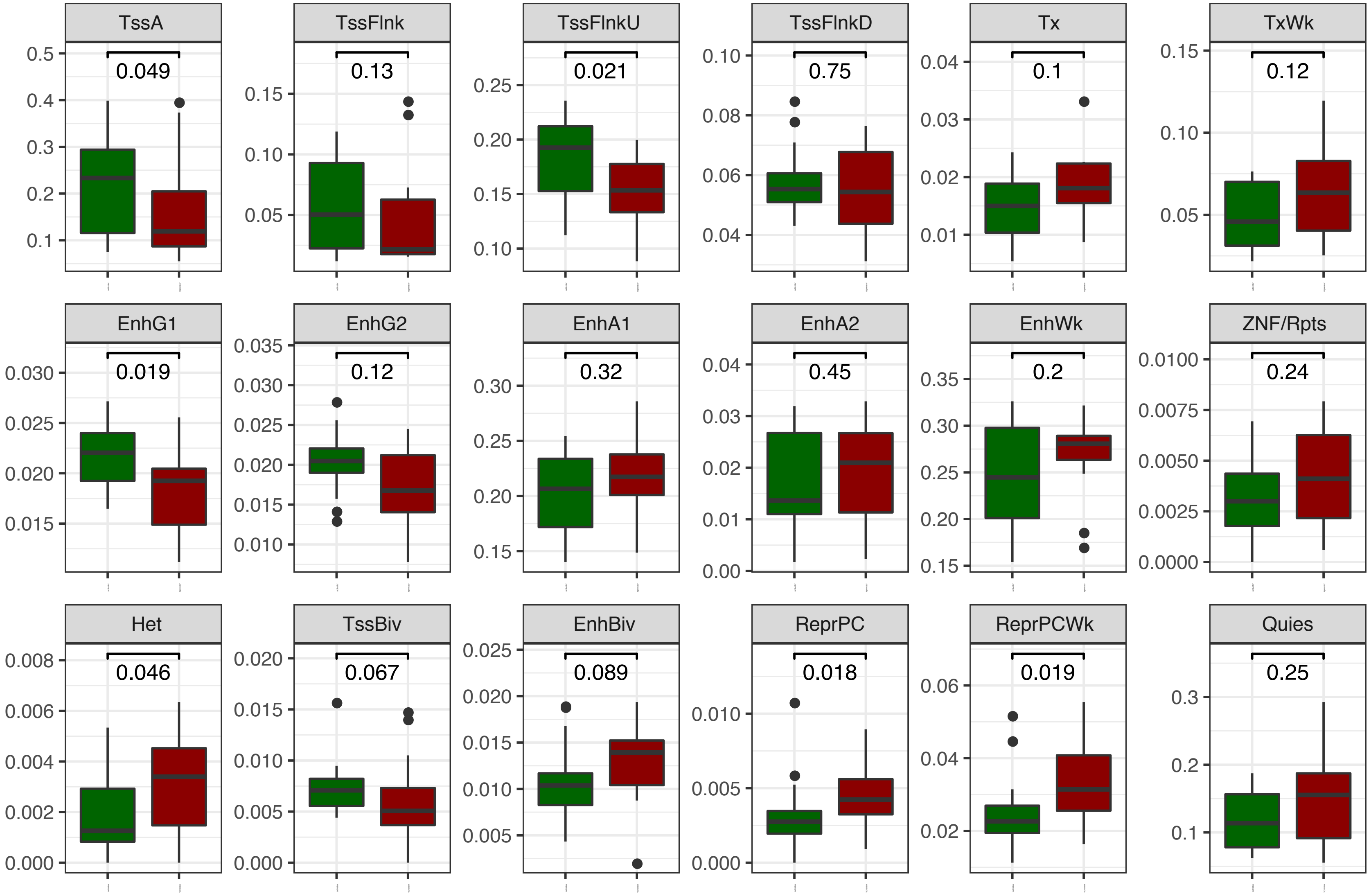

TssA
0.049
TssFlnk
0.13
TssFlnkU
0.021
TssFlnkD
0.75
Tx
0.1
TxWk
0.12
EnhG1
0.019
EnhG2
0.12
EnhA1
0.32
EnhA2
0.45
EnhWk
0.2
ZNF/Rpts
0.24
Het
0.046
TssBiv
0.067
EnhBiv
0.089
ReprPC
0.018
ReprPCWk
0.019
Quies
0.25

**Supplementary Figure 16.** Additional results and alternative visualizations for the GMP-MPP dataset. (A) Volcano plot in analogy to Figure 2a. (B) Raw p-value histogram for the analytical approach. (C) Comparison of the empirical and analytical approach. While the empirical approach uses permutations to estimate significance, the analytical one estimates significance using the methods and formulas as described in the methods part. The y-axis shows the result for the analytical approach, the x-axis for the empirical one with respect to differential TF activity (left) and ranks of adjusted p-values (right). Note that we used the “first” method to break ties, because the empirical approach may produce p-values that are exactly 0 for a particular set of TFs, all of which then have a tied rank.

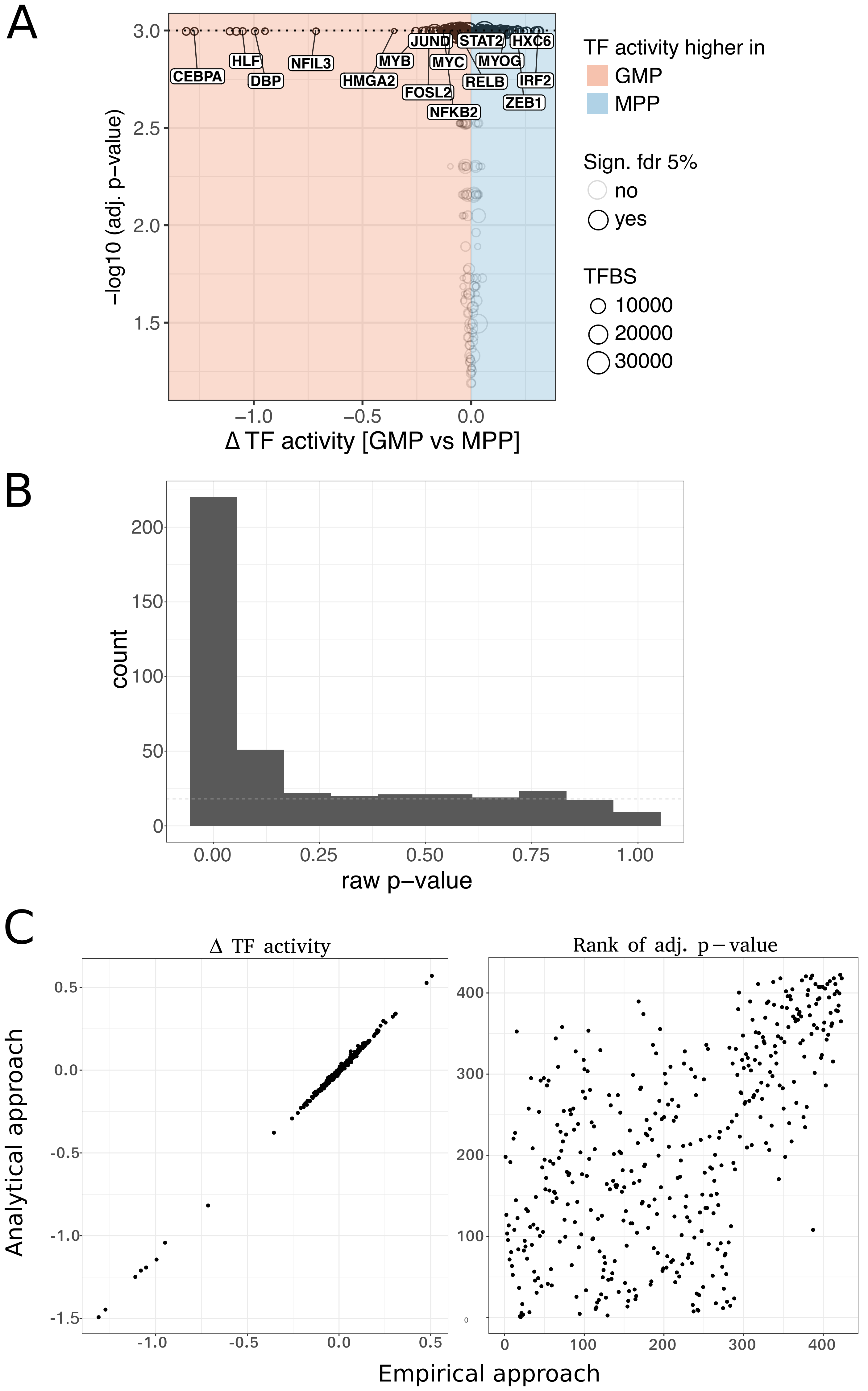
A
CEBPA
HLF
DBP
NFIL3
MYB
HMGA2
JUND
MYC
FOSL2
NFKB2
STAT2
MYOG
RELB
HXC6
IRF2
ZEB1
−log10 (adj. p−value)
Δ TF activity [GMP vs MPP]
TF activity higher in
GMP
MPP
Sign. fdr 5%
no
yes
TFBS
10000
20000
30000
B
count
raw p−value
C
Δ TF activity
Rank of adj. p − value
Analytical approach
Empirical approach

**Supplementary Figure 17.** Cluster heatmap for the footprints (left) and summary cluster footprints (right) for the GMP/MPP dataset. This Figure is an extension of Fig. 6b and shows all clusters and the clustering tree. The heatmap shows TF footprint values for all significant TFs from *diffTF* that were also significantly differentially expressed (adj. p-value $< 0.05$ for both), based on the normalised Tn5 insertions for each bp (+/-100bp from motif center). Colors represent footprint strength, while white denotes the value of the genomic background in the consensus peakset. Clusters were defined using hierarchical clustering with the *ward.D2* method. For the cluster summary footprints at the left, we divided each footprint value by the mean value of each cluster in order to highlight the differences in the surrounding chromatin structure and colored them according to their hypothesized footprint type into activators (green) and repressors (red). The direction of TF expression and TF activity is in analogy to what is described in the text. “Direction” denotes whether expression and TF activity have the same or opposite sign.

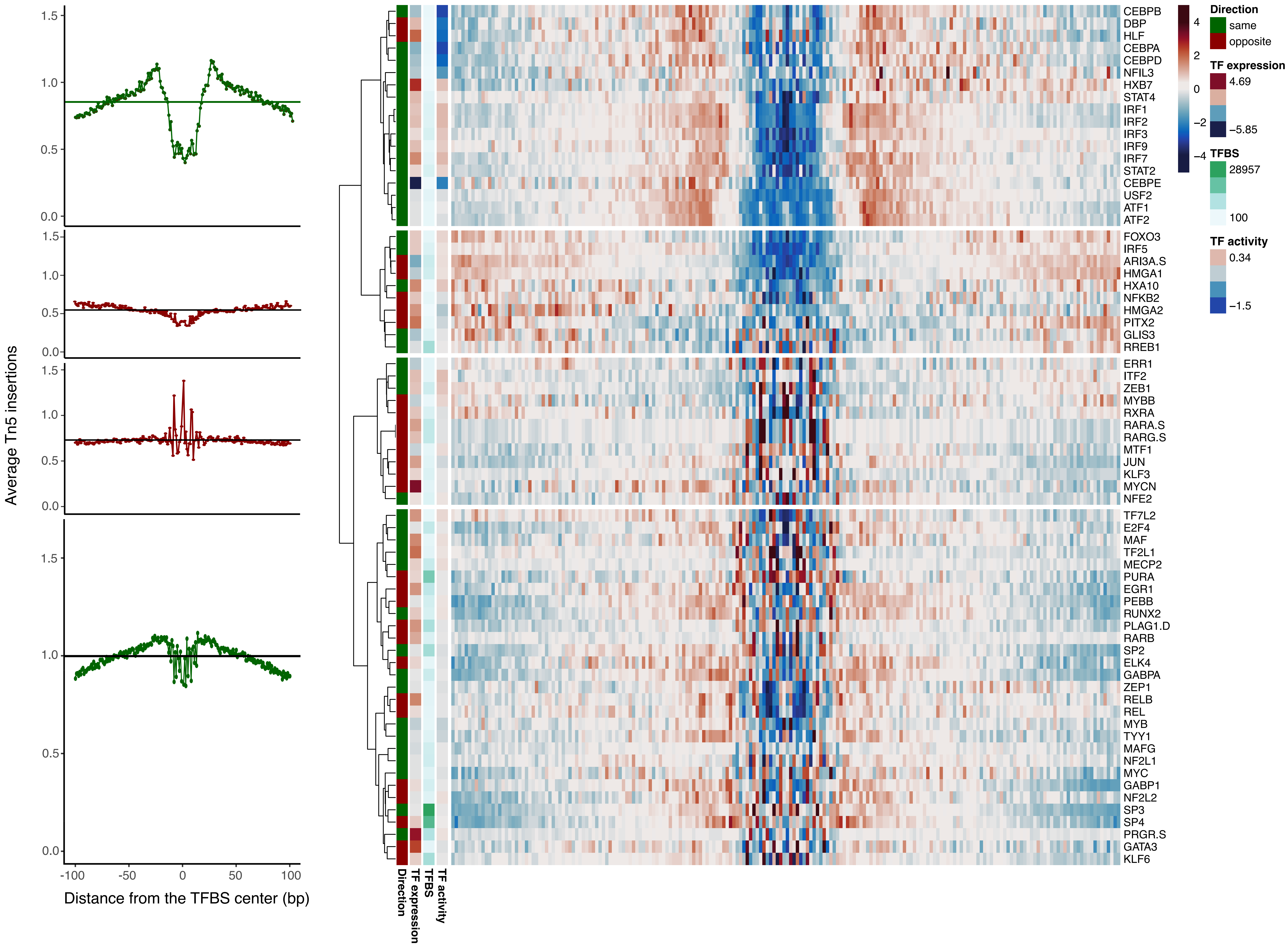

Average Tn5 insertions
Distance from the TFBS center (bp)
Direction
TF expression
TFBS
TF activity
Direction
same
opposite
TF expression
4.69
−5.85
TFBS
28957
100
TF activity
0.34
−1.5
CEBPB
DBP
HLF
CEBPA
CEBPD
NFIL3
HXB7
STAT4
IRF1
IRF2
IRF3
IRF9
IRF7
STAT2
CEBPE
USF2
ATF1
ATF2
FOXO3
IRF5
ARI3A.S
HMGA1
HXA10
NFKB2
HMGA2
PITX2
GLIS3
RREB1
ERR1
ITF2
ZEB1
MYBB
RXRA
RARA.S
RARG.S
MTF1
JUN
KLF3
MYCN
NFE2
TF7L2
E2F4
MAF
TF2L1
MECP2
PURA
EGR1
PEBB
RUNX2
PLAG1.D
RARB
SP2
ELK4
GABPA
ZEP1
RELB
REL
MYB
TYY1
MAFG
NF2L1
MYC
GABP1
NF2L2
SP3
SP4
PRGR.S
GATA3
KLF6

**Supplementary Figure 18.** Comparison of *diffTF* and *HOMER* for M-CLL (A) and U-CLL (B). Each point represents one TF. The x-axis shows the differential TF activity from *diffTF*, while the *HOMER* enrichment (in %) is shown on the y-axis. Colors represent significance (FDR < 0.1): white - not significant in either analysis; light green and light blue - significant only for the analysis from the x-axis or y-axis, respectively; purple - significant for both analyses. Spearman correlation was computed only for TFs with FDR < 0.1 for the adjusted p-values from the *HOMER* enrichment using Benjamini-Hochberg. 0 and 32 motifs were enriched in *HOMER* in (A) and (B), respectively. In (B), a linear model is shown as a black line.

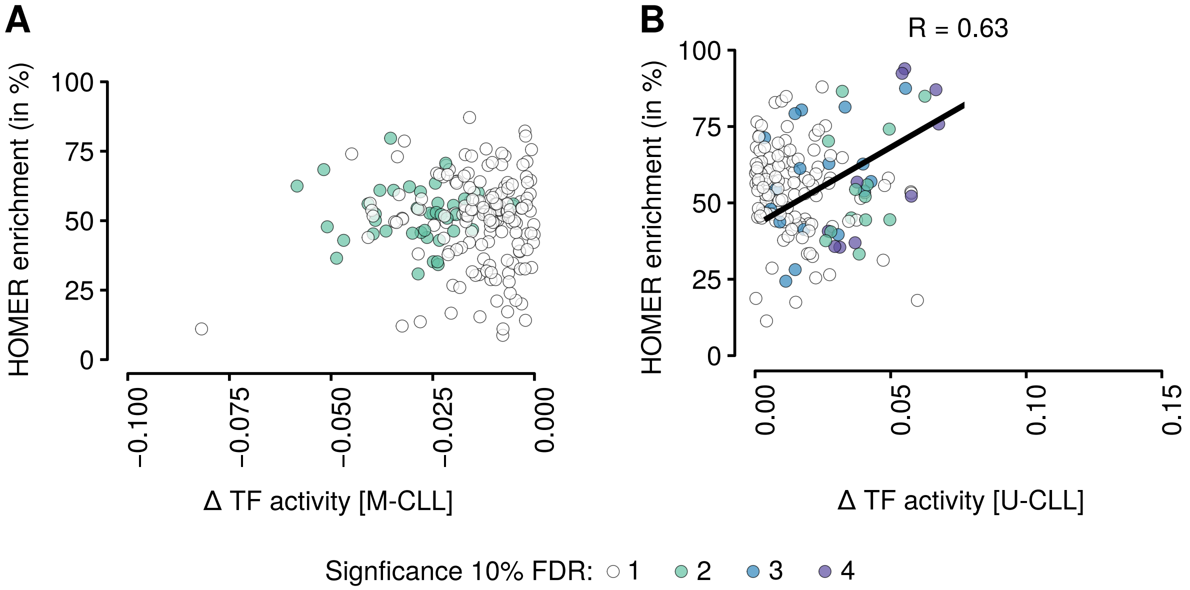

A
B
R = 0.63
HOMER enrichment (in %)
100
75
50
25
0
−0.100
−0.075
−0.050
−0.025
0.000
0.00
0.05
0.10
0.15
Δ TF activity [M-CLL]
Δ TF activity [U-CLL]
Signficance 10% FDR: 1 2 3 4

**Supplementary Figure 19.** Comparison plots for *diffTF* vs. *chromVAR*. A-B: Both plots show the *diffTF* TF activity on the x-axis and the *chromVAR* deviation (A) and deviation score (B) on the y-axis, as measured by the difference of the means between the two conditions U-CLL and M-CLL. Only the 370 expressed TFs are included. Pearson/Spearman correlation for A: 0.8/0.79, B: 0.81/0.77. The three TFs that are mentioned in C, D and F are labeled in A. C-D: Correlation of log2 fold-changes between U-CLL and M-CLL from peaks versus individual TFBS for two selected TFs. Note that for each peak, the TFBS per TF with the highest average read count across all samples was selected (see *diffTF* workflow). (C) TF with the lowest correlation among all expressed TFs from the second quadrant (NFIL3, Pearson 0.84, 290 TFBS). (D) TF with the highest correlation among the set of significant TFs according to *diffTF* (IRF2, Pearson 0.95, 4362 TFBS). E: Summary of TF-specific differences between mean log2 fold-changes of peaks vs. TFBS (see C and D) across all TF. F: Density of *chromVAR* deviations for CUX1, the summary score of which turns positive when computed as the difference between the mean deviation per group (0.0001028 - -0.0000952 = 0.000198) due to one outlier sample in M-CLL.

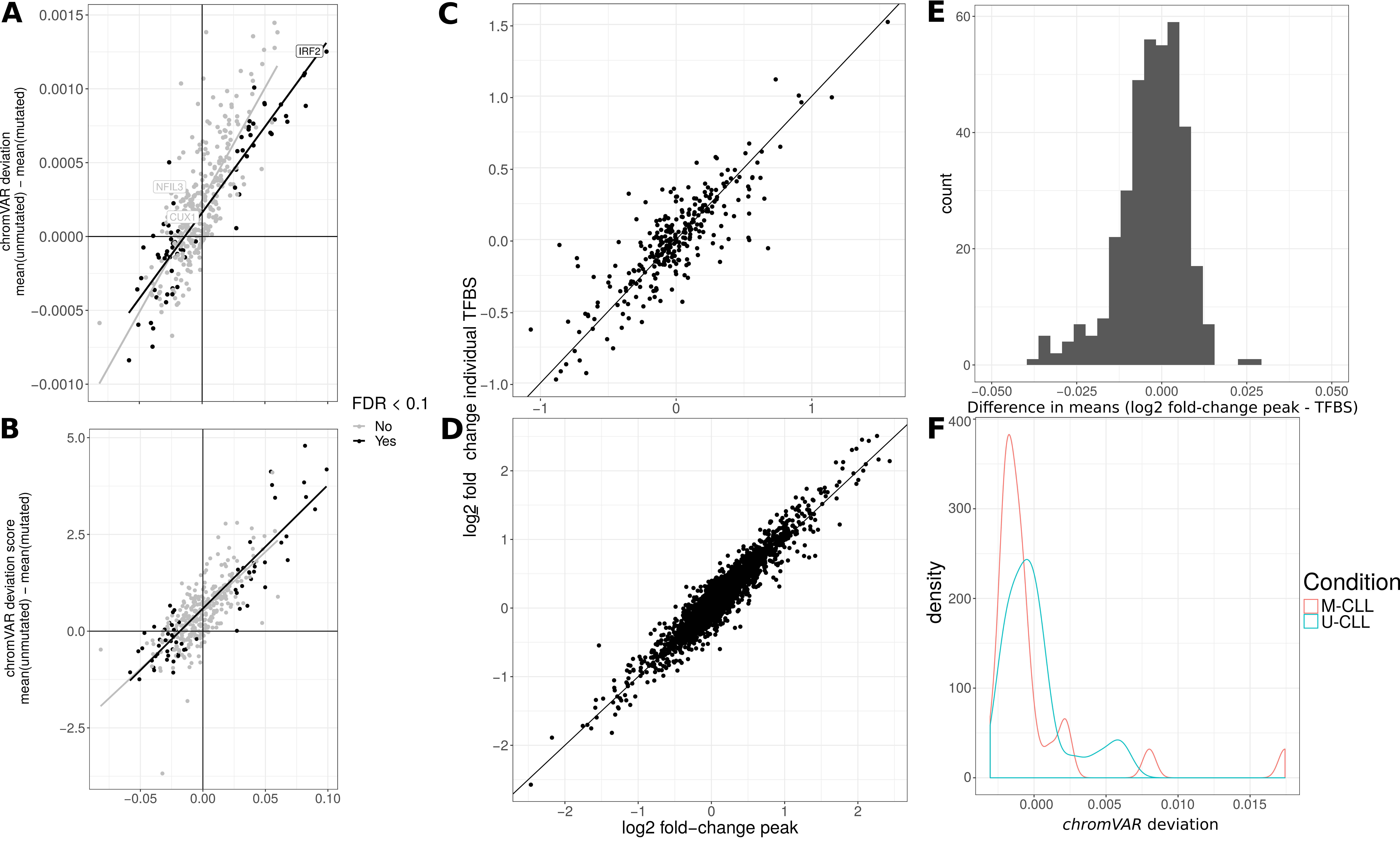
A
B
C
D
E
F
IRF2
NFIL3
CUX1
chromVAR deviation
mean(unmutated) − mean(mutated)
chromVAR deviation score
mean(unmutated) − mean(mutated)
FDR < 0.1
No
Yes
log2 fold change individual TFBS
log2 fold−change peak
count
Difference in means (log2 fold-change peak - TFBS)
density
chromVAR deviation
Condition
M-CLL
U-CLL

### Supplementary Tables:

**Supplementary Table 1.** Final results of the *diffTF* pipeline for the CLL data. The columns contain the TF (*TF*), the number of TF binding sites (*TFBS*), the differential TF activity (*differential_TFActivity*), the raw and adjusted p-values (*pvalue* and *pvalueAdj*, respectively) the TF class based on the correlation of RNA-Seq and ATAC-Seq data (*classification*) as well as the median Pearson correlation of TF-containing peaks and TF expression (*AR correlation*).

**Supplementary Table 2.** List of all significantly differentially active TFs and their annotated condition-specific CLL associations. Data for TF families and DNA binding domains were obtained from *GeneCards* and http://humantfs.ccbr.utoronto.ca, respectively. For the TF annotation we assume that if the one TF is annotated in the literature with the specific condition (M-CLL or U-CLL) we counted the whole functional family as annotated if PWMs are closely related in terms of motif similarity.

**Supplementary Table 3.** Metadata for the CLL dataset and list of TF that we used in all analyses, including a list of TFs overlapping with *ReMap 2015*.

**Supplementary Table 4.** Final results of the *diffTF* pipeline for the MPP/GMP dataset. The structure is identical to the table in Supplementary Table 1.

**Supplementary Table 5.** Qualitative comparison of *diffTF* with similar tools. Comparison and distinction of *diffTF* with regard to existing tools for a similar purpose. Green, yellow and red: Feature fully, partially or not supported/implemented/available, respectively. Gray indicates non-applicability.
